# Derivation of elephant induced pluripotent stem cells

**DOI:** 10.1101/2024.03.05.583606

**Authors:** Evan Appleton, Kyunghee Hong, Cristina Rodríguez-Caycedo, Yoshiaki Tanaka, Asaf Ashkenazy-Titelman, Ketaki Bhide, Cody Rasmussen-Ivey, Xochitl Ambriz-Peña, Nataly Korover, Hao Bai, Ana Quieroz, Jorgen Nelson, Grishma Rathod, Gregory Knox, Miles Morgan, Nandini Malviya, Kairui Zhang, Brody McNutt, James Kehler, Amanda Kowalczyk, Austin Bow, Bryan McLendon, Brandi Cantarel, Matt James, Christopher E. Mason, Charles Gray, Karl R. Koehler, Virginia Pearson, Ben Lamm, George Church, Eriona Hysolli

## Abstract

The crisis of biodiversity loss in the anthropogenic era requires new tools for studying non-model organisms. Elephants, for example, are both an endangered species and excellent models studying complex phenotypes like size, social behavior, and longevity, but they remain severely understudied. Here we report the first derivation of elephant (*Elephas maximus*) induced pluripotent stem cells (emiPSCs) achieved via a two-step process of chemical-media induction and colony selection, followed by overexpression of elephant transcription factors *OCT4, SOX2, KLF4, MYC* ± *NANOG* and *LIN28A*, and modulation of the *TP53* pathway. Since the seminal discovery of reprogramming by Shinya Yamanaka, iPSCs from many species including the functionally extinct northern white rhinocerous have been reported, but emiPSCs have remained elusive. While for multiple species the reprogramming protocol was adopted with little changes compared to model organisms like mouse and human, our emiPSC protocol requires a longer timeline and inhibition of *TP53* expansion genes that are hypothesized to confer unique cancer resistance in elephants. iPSCs unlock tremendous potential to explore cell fate determination, cell and tissue development, cell therapies, drug screening, disease modeling, cancer development, gametogenesis and beyond to further our understanding of this iconic megafauna. This study opens new frontiers in advanced non-model organism cellular models for genetic rescue and conservation.

## 1 Introduction

Over the past 10,000 years, humans have had a profound impact on the world’s ecosystems. Humans have completely changed the ecology of nearly every ecosystem in the world and were potentially at least partially responsible for the extinction of more than two thirds of the world’s land megafauna [1, 2]. Today we see that many of the world’s remaining megafauna are also edging towards extinction [3]. Among these are the world’s largest remaining land animals - the elephants. In recent centuries, international efforts have formed to conserve and preserve these megafauna.

Until the last decade, a majority of these efforts were operational, but recently multiple groups have considered using biotechnology as a conservation tool [4]. Moreover, biotechnology is now being developed as a ‘de-extinction’ solution to bring back species that have gone extinct and left gaps in their respective ecosystems, like the woolly mammoth [5].

A key component of these conservation and de-extinction workflows is the generation of induced pluripotent stem cells (iPSCs). iPSCs share the molecular and phenotypic features of embryonic stem cells (ESCs) and can, in principle, be converted into any other type of cell in an organism’s body — including embryonic cells needed for de-extinction and conservation. iPSCs were first derived from somatic cells from mouse [6], and, shortly after, humans [7]. At present, mouse and human iPSCs have been studied and engineered extensively, but iPSCs from other species lag behind. iPSCs have been demonstrated for many animals such as cows, pigs, goats, sheep, horses, marmosets, dogs, rabbits, rhinoceroses, naked mole rats, and bats [8, 9, 10, 11, 12], however, for many animal species such as elephants and whales, iPSC-creation has remained elusive. To this end, scientists have begun biobanking tissue samples and cell lines from many endangered species in the hopes that iPSCs may someday be derived.

With iPSCs from each species, doors are opened to understanding their biology and engineering their survival or re-birth. In recent years, stem cell biologists have engineered cell-fate with stem cells to differentiate them into many different types of cells for therapeutic or diagnostic purposes [13, 14, 15, 16, 17]. By creating stem cells and diverse differentiated cell type populations from stem cells, biologists can explore the features of specific cell types in vitro and identify key features unique to that species. Stem cell biologists have also created entire organisms from iPSCs using 4N-complementation [18], chimeric embryos from stem cells originating from two different species [19], and embryo models from iPSCs [20, 21]. These advances make the idea of species conservation and de-extinction ever more feasible, but require the creation of iPSCs from each species and the re-solving of frame solutions demonstrated from others.

Beyond their ecological relevance, elephants have very interesting biological capabilities — they are highly intelligent [22], exhibit interesting molecular aging features common to humans [23, 24], and rarely get cancer [25]. Their resistance to cancer has drawn significant attention in recent years — given their huge size, elephants should get cancer at a much high rate than they do, a paradox called Peto’s paradox [26]. While this paradox has been studied in elephants before [27], the way in which elephants solve this paradox has remained unsolved. Creation of elephant stem cells could shed light into solving this problem given the overlap of many core gene networks involved in stem cells and cancer cells. Moreover, Asian elephants have 29 *TP53* and 8 *LIF* polymorphic gene copies in the genome[28, 29], both of which are established key genes in both pluripotency and cancer [30].

Here, we describe the generation of *Elephas maximus* induced pluripotent stem cells (emiPSCs) with our multidimensional reprogramming protocol, a chemical-based approach followed by core reprogramming transcription factor overexpression. Next, we studied the key pluripotency molecular features of emiPSCs and characterized their ability to differentiate. Via comparative genomics we were able to establish a comparison to stem cells of other mammals and showed that their features align closest with rhino and cattle, two other large living land mammals.

## 2 Results

### 2.1 Asian elephant reprogramming

After a comprehensive screen of more traditional reprogramming approaches failed, elephant induced pluripotent stem cells were generated using chemical-based pluripotency media with selected colony expansion, followed by overexpression of key pluripotency transcription factors OSKM(LN) and oncogene SV40LT and/or short hairpin RNAs (shRNAs) against elephant *TP53*. Multiple attempts with current standard reprogramming methods were tried, and failed, and resulted in no, or incomplete, reprogramming. First, we attempted to reprogram *Elephas maximus* cells via the use of over-expression of standard ‘Yamanaka reprogramming factors’: *OCT4, SOX2, KLF4*, and *MYC* (OSKM) in different vector formats. This included transgene expression via episomal [31], lenti-virus [32], Sendai-virus [33], and PiggyBac [34] in a variety of different combinations with transcription factor (TF) sequences from either mouse or human. These transgene expression methods were also tested in tandem with the over-expression of shRNAs that target TP53, SV40 T-antigen, *NANOG*, and/or *LIN28A*. While some of these methods yielded some potential morphological differences from primary cells at different parts of the process, ultimately all of these types of attempts failed as a result of either cell death, senescence, or no observed morphological changes compared to starting primary parental lines (**Supplementary Figure 1-9**).

In addition to transgene-dependent methods, we also explored a growing set of chemical reprogramming methods that do not require the use of transgenes. We tested a set of published cocktail sets [35, 36, 37] that we further customized by testing multiple variations of chemical cocktails. These approaches resulted mostly in cell death, senescence, or no observed change, but in a small set of experiments also appeared to yield encouraging morphological changes (**Supplementary Figure 1-9**).

It was through integration of these two types of approaches that resulted in the first generation of *Elephas maximus* induced pluripotent stem cells (emiPSCs). More specifically, we derived emiPSCs by first partially reprogramming primary endothelial cells (emECs) to an intermediate ‘pre-iPSC’ state (emPRC) with a derivative chemical reprogramming protocol based on prior work [35], and then using elephant-specific transgene (Elephant AA sequence homology percentages to human Yamanaka factors are: *OCT4* - 88.4%, *SOX2* - 98.1%, *KLF4* - 88.6%, *MYC* - 91.1%) over-expression to complete the reprogramming process (**Figure 1a**). While encouraging changes to morphology compared to primary cells was observed through chemical treatment alone (**Figure 1b,c**, **Supplementary Figure 4e**), we observed a more canonical stem cell morphology and growth rate only after introduction of pluripotency transgenes (**Figure 1d**, **Supplementary Figure 4f**). However, even more mature iPSCs were difficult to establish, as mouse and human reprogramming transcription factors were not sufficient. Furthermore, no meaningful change was observed in either growth, morphology or molecular composition until we modulated the *TP53* expression. We achieved the latter by over-expression of SV40 Large T-antigen or an shRNA that targeted RNA *only* from *TP53* retrogenes and not the full-length *TP53* gene (**Supplementary Figure 5**). Interestingly, in the reverse order, when primary cells were treated to induce transgenes before the treatment with the chemical cocktail, cells quickly senesced and underwent apoptosis in all tested conditions.

**Figure 1:**
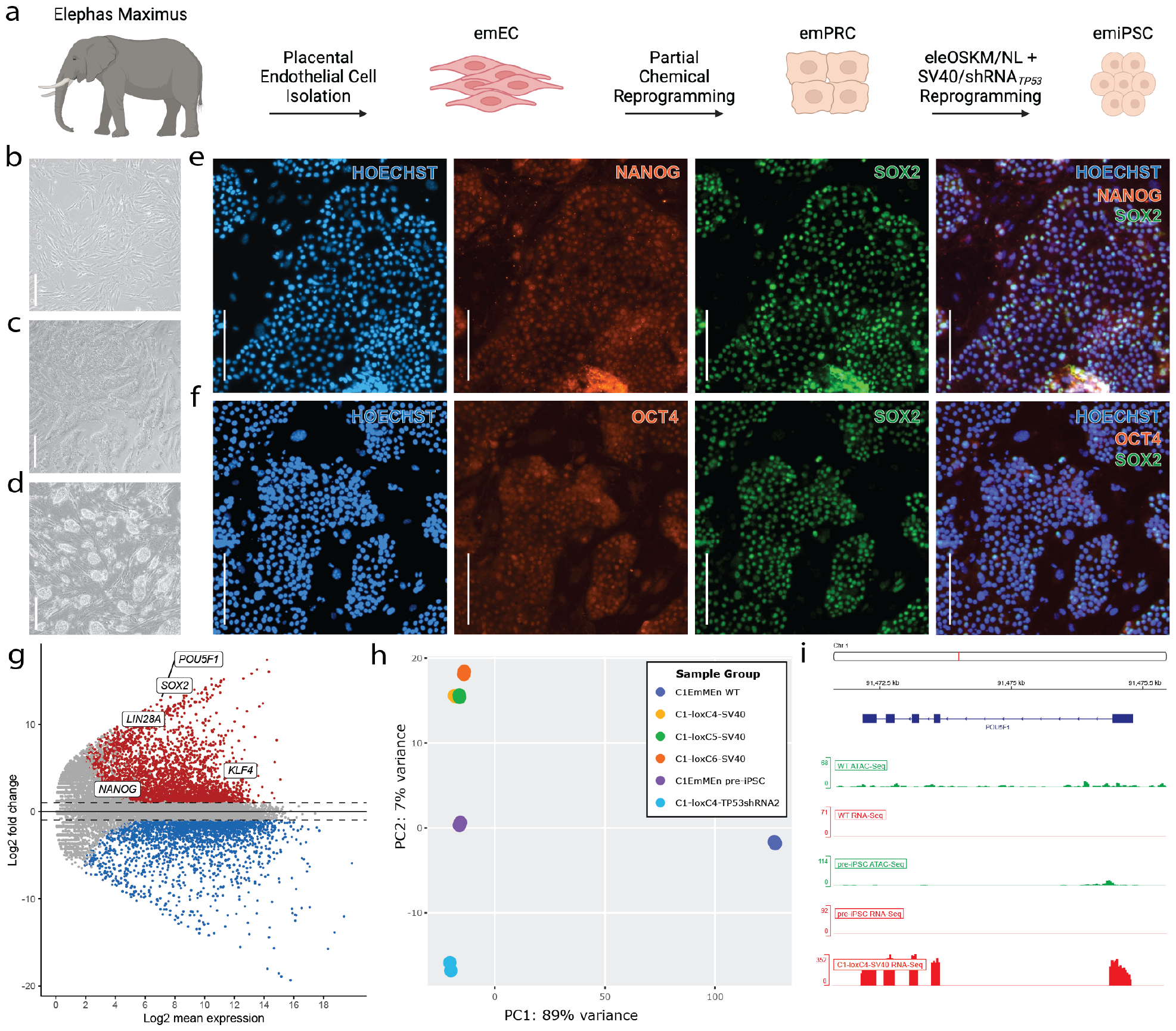
Derivation of *Elephas maximus* induced pluripotent stem cells. **a**. Illustration of the reprogramming strategy involved. emEC - *Elephas maximus* endothelial or epithelial cells; emPRC - *Elephas maximus* partially reprogrammed cells; eleOSKM/NL - *Loxodonta africana POU5F1, SOX2, KLF4, CMYC, LIN28A*, and *NANOG* transgenes; shRNA_*T P* 53_ - shRNAs targeting *TP53* and its retrogenes. **b.-f**. All scale bars = 200*μ*M, magnification = 10X **b**. Brightfield (BF) image of *Elephas maximus* primary endothelial cells. (emEC) **c**. BF image of partially reprogrammed (pre-iPSC) cells (emPRC) **d**. BF image of *Elephas maximus* induced-pluripotent stem cells. (emiSPCs) **e**. Immuno-fluorescent detection of *NANOG* (Texas Red) / *SOX2* (AF488) / HOECHST separately and merged. **f**. Immuno-fluorescent detection of *OCT4* (Texas Red) / *SOX2* (AF488) / HOECHST separately and merged. **g**. MA plot of RNA-seq data illustrating the transcriptional differences between elephant endothelial cells (emEC) and pluripotent stem cells (emiPSC). Key canonical pluripotency genes are labeled. **h**. PCA plot comparing emECs (C1EmMEn WT), emPCRs (C1EmMEn pre-iPSC), and emiPSCs (C1-loxC4/5/6-SV40 and C1-loxC4-TP53shRNA2). **i**. RNA-seq and ATAC-seq signals of *OCT4* (*POU5F1*) for emECs, emPRCs, and emiPSCs. Shown are peak calls of one representative sample.

Upon establishing a more canonical stem cell morphology, we pursued a full stem cell characterization. We demonstrated that our cells express core pluripotency proteins via immunofluorescence (IF) (**Figure 1e,f, Supplementary Figure 10**). Since elephants often have a considerably diverged amino acid sequence compared to human, mouse, and other studied mammal sequences, we contracted the creation and validation of elephant amino-acid-sequence-specific antibodies for IF staining. In some cases where the selected epitope of the custom-designed antibody overlapped perfectly with antibodies targeting the same region, we substituted commercial antibodies after testing positive controls in human iPSCs. Next, we confirmed via RNA-seq that these core genes and others are indeed present in four derivative emiPSC lines (**Figure 1g**). Interestingly, upon a further examination of the transcriptome of our emiPSC lines, emPRCs, and emECs, we saw nearly 90% of the transcriptome differences were accomplished with the chemical cocktail alone (**Figure 1h**). Finally, we wanted to examine *OCT4* (*POU5F1*) epigenetic and transcriptomic read-mapping to determine if perhaps the chemical pre-treatment yielded increased regions of open chromatin upstream of the *OCT4* locus (**Figure 1i**). It appears that this indeed was not the case, confirming that the chemical cocktail does a lot of work bears the brunt of modulating the overall epigenetic landscape of these cells (**Supplementary Figure 8,9**), but it is not sufficient to modulate chromatin accessibility upstream of *OCT4*.

While we successfully conducted a multitude of molecular and functional tests for our emiPSCs, interestingly, we did not observe a significant upregulation of *NANOG* expression in our emiPSCs. This expression pattern is unusual for iPSCs of species studied to date. To address this point, we performed another round of nucleofection with pluripotency genes to attempt to boost *NANOG* expression in these cells. We added extra copies of both elephant Yamanaka factors (OSKM), and *NANOG* individually (**Supplementary Figure 11**). In order to show increased expression of endogenous *NANOG* (not simply transgene expression), we performed RT-qPCR with primers that bridge intron-exon gaps of mRNA in *NANOG* and the coding sequence - 3’ UTR region of *SOX2*, otherwise not found in transgenes. Our results demonstrate that we are able to boost endogenous *NANOG* (and endogenous *SOX2*) expression in our emiPSC lines.

After establishing a boost to endogenous expression of *NANOG* and *SOX2*, we asked whether our cells could withstand the withdrawal of transgene over-expression driven by doxycycline (DOX). We withdrew DOX for 10 days and observed via RNAseq that while the expression of *OCT4* and *SOX2* decreased, they remained significantly upregulated compared to parental cells (**Supplementary Figure 12a**). Furthermore, the expression profile of other core pluripotency markers and cell-cycle regulatory markers remained mostly unchanged (**Supplementary Figure 12b-d**), thus we withdrew DOX from downstream differentiation studies.

### 2.2 Asian elephant pluripotent stem cells

Our emiPSCs express multiple naïve pluripotency markers, and are capable of differentiation into three germ layers as assessed by embryoid body differentiation and teratoma formation. Once the expression of core pluripotency markers was established, we wanted to evaluate the basic growth and chromosomal normality of the cells. To do this end, we performed a simple nuclei isolation, lysis, and chromosomal counting methodology to confirm that our cells indeed had 56/56 of the expected chromosome number (**Figure 2a**). Next, we measured the doubling time of our set of emiPSCs on mouse embryonic feeder cells (MEFs) (**Figure 2b**) and determined that the doubling time for each line was approximately slightly over 2 days, which seemed reasonable given the 2-year elephant gestation period.

**Figure 2:**
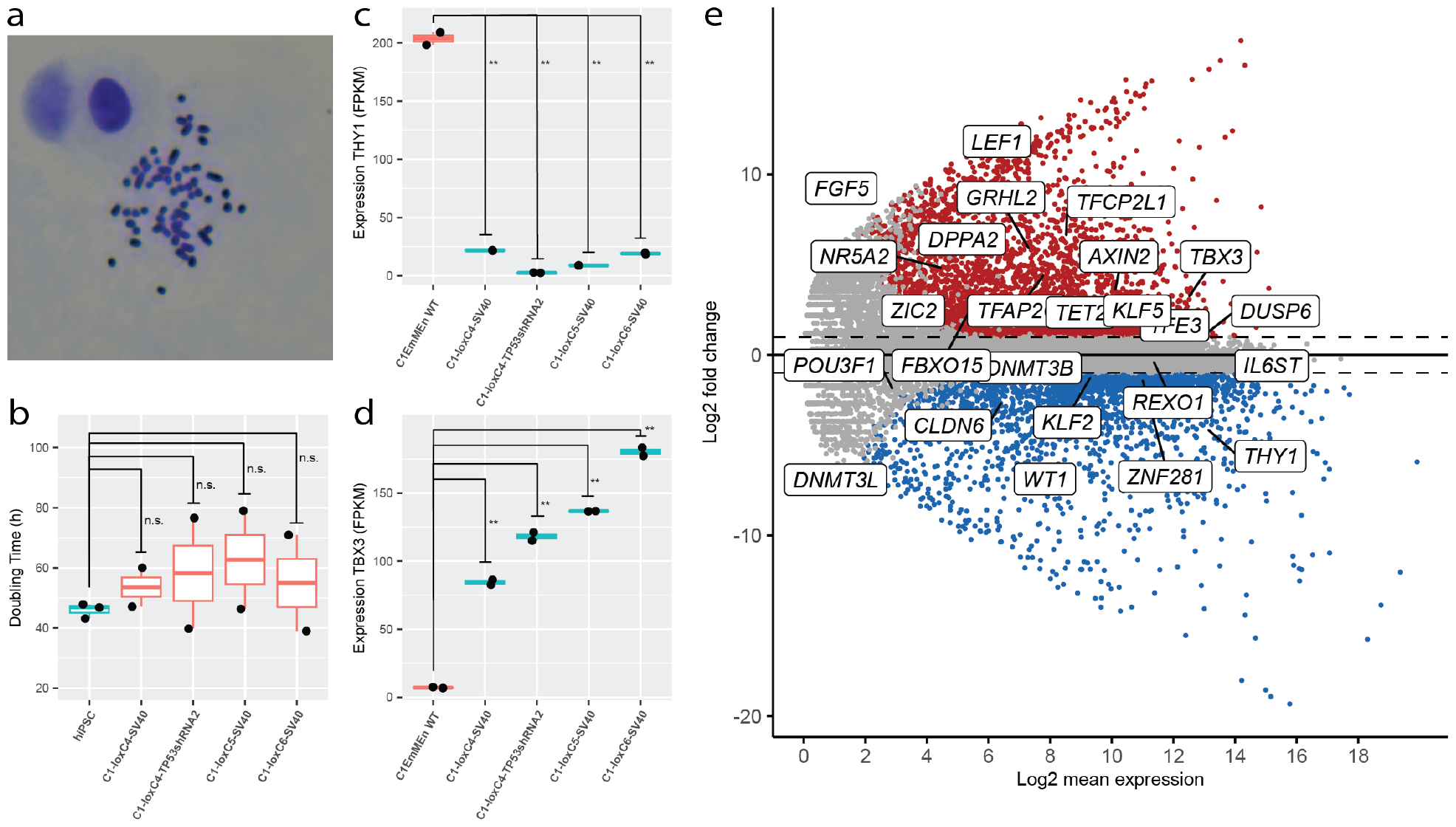
Characteristics of *Elephas maximus* induced pluripotent stem cells. **a**. Karyotyping staining of emiPSC nuclei show 56/56 chromosomes **b**. Doubling time for emiPSC cell lines and hiPSCs as originally derived in 2007. Statistical tests show all emiPSCs are not statistically significantly slower growing that originally derived hiPSCs. **c**. Expression of a canonical primed pluripotency markers *THY1* and a **d**. canonical naive pluripotency marker *TBX3*. All emiPSC lines significantly higher expression than WT controls (p < 0.01). **e**. MA plot showing a large set of canonical naive and primed pluripotency markers for C1-loxC4-SV40. A vast majority of naive markers are up-regulated and most primed markers are down-regulated.

Next, we wanted to examine gene expression of genes canonically expressed in primed versus naive iPSCs and determine whether our cells were more primed or naïve. We observed that our cells exhibited morphology more generally associated with naïve cells in other species and have negligible spontaneous differentiation observed (**Figure 1d**). We focused on the expression of canonical markers that could shed more light on this observation. For primed marker *THY1*, we find very highly significant down-regulation across emiPSC lines (**Figure 2c**), and highly significant up-regulation of *TBX3* (**Figure 2d**). However, across the board with most naive cell markers, we see significant upregulation of emiPSCs compared to emECs (**Figure 2e**). There are indeed some canonical primed markers seen significantly upregulated (*ZIC2* and *GRHL2*), but it is known that primed and naive pluripotency markers vary somewhat across species.

Since our emiPSC lines demonstrated expected molecular and phenotypic qualities of stem cells and a normal karyotype, we next probed their differentiation potential. First, we performed an embryoid body (EB) formation assay. We demonstrate that all our cell lines form EBs in 5-7 days, and that they contain cells that express early differentiation markers from each of the three germ layers - *PAX6* (ectoderm), *GATA4* (mesoderm), and *FOXA2* (endoderm) (**Figure 3a,b**, **Supplementary Figure 13**). Unlike emPRC, emiPSCs consistently formed EBs, thus we performed RNA sequencing on these populations to look for the presence of additional early differentiation markers. We highlight the expression of endoderm markers *PRDM1, FOXA2* and *EOMES*; ectoderm markers *PAX6, PAPLN* and *POU4F1*, and mesoderm markers *CDX2, GATA4* and *HAND1* (**Figure 3c**). We also show that the upregulation of these early differentiation markers in EBs increases over time to further confirm these trends (**Supplementary Figure 14**). Many core differentiation markers are upregulated compared to non-reprogrammed parental cells, but we acknowledge variations compared to other species that need further examination.

**Figure 3:**
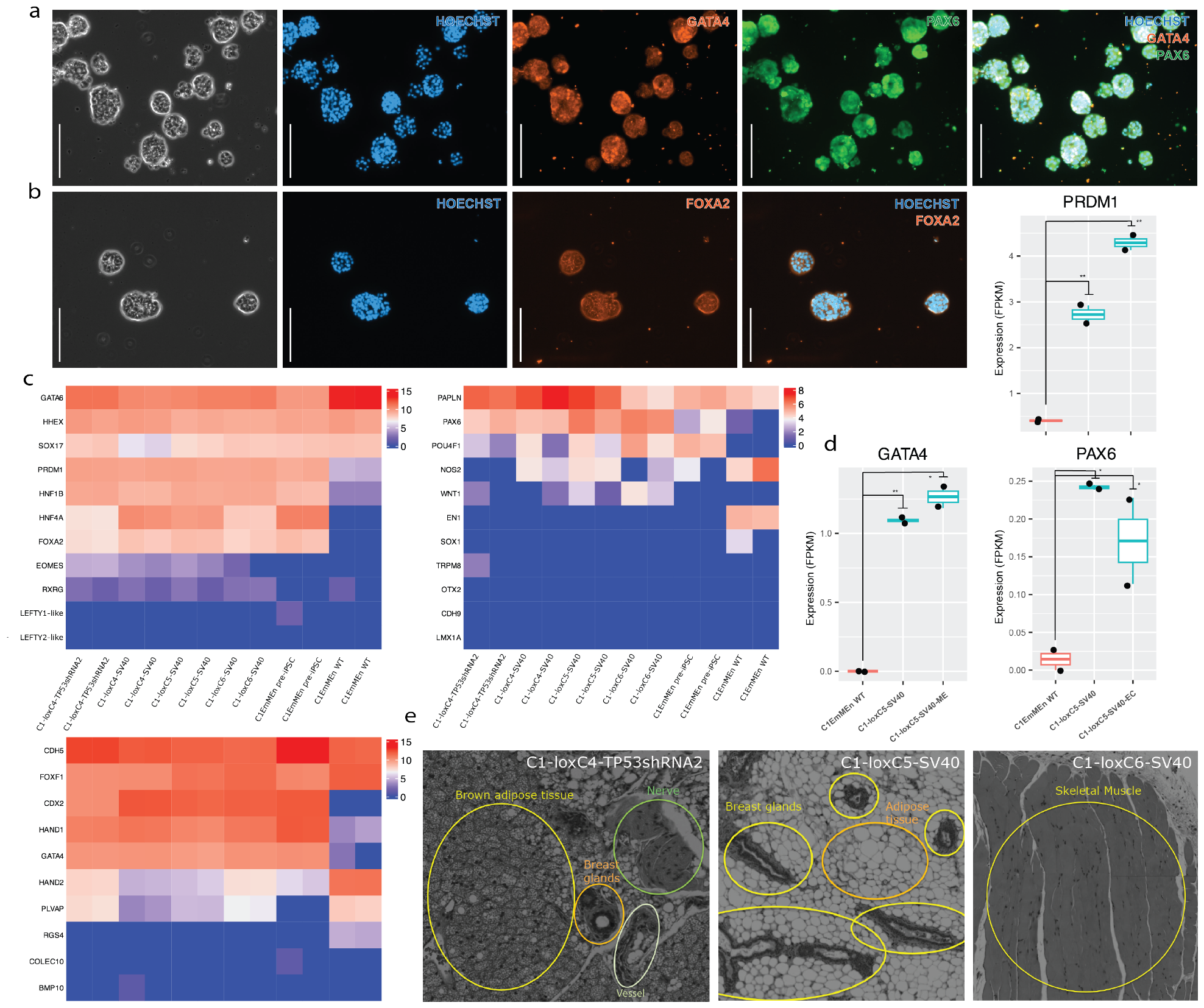
Differentiation potential of *Elephas maximus* induced pluripotent stem cells. **a.-b**. All scale bars = 200uM, magnification = 10X **a**. Immunofluorescence (IF) microscopy images of embryoid bodies (EBs) formed by emiPSC line C1-loxC5-SV40 co-stained with antibodies detecting the expression of lineage-specific markers *PAX6* (ectoderm) and *GATA4* (mesoderm). **b**. IF microscopy images of EBs formed by emiPSC line C1-loxC5-SV40 stained with an antibody detecting the expression of lineage-specific markers *FOXA2* (endoderm). **c**. Representation of canonical germ-layer differentiation marker genes in EBs formed from each emiPSC line, emPRCs, and emECs. (LEFT) Endoderm markers, (RIGHT) ectoderm markers, and (BOTTOM) mesoderm markers. **d**. Expression of early differentiation markers *PAX6* (ectoderm), *GATA4* (mesoderm), and *PRDM1* (endoderm) following tri-lineage differentiation in emiPSC line C1-loxC5-SV40. Statistical t tests show ** = p < 0.01 and * = p < 0.05. **e**. Microscopic images of hematoxylin-eosin-stained sections of tumor tissue after injection of emiPSCs (to form teratoma) into the upper hind-leg of immuno-compromised mice. Teratomas formed from each emiPSC line resulted in different cell types being developed (marked by color in sub-panels).

After confirming the presence of many of the expected early differentiation markers in EBs, we performed tri-lineage differentiation assays to verify the presence of marker genes in their respective lineages. We observed up-regulation of early differentiation markers in these differentiated cells compared to both their originating emiPSC line and emECs via RNAseq (**Figure 3d**, **Supplementary Figure 15**).

Given that our iPSC lines could form EBs and differentiate into the early three germ layers, we tested whether these lines could form teratomas. While this assay can be confounding and its presence or absence in datasets is researcher-dependent most likely because it is generally hard to standardize [38], we further pursued it as an additional assay to probe the differentiation potential of our cells. To this end, we injected emiPSCs into hind-leg of immuno-compromised mice and observed growth for 4-6 weeks. We could detect formation of teratomas, and furthermore, multiple tissue types were observed in the teratomas formed from each cell line (**Figure 3e**, **Supplementary Figure 16**). Interestingly, while the molecular differences between our emiPSC lines was relatively minor, we appeared to see more variety of cell types in the cell line that contained an shRNA against *TP53* retrogenes, than the lines which required SV40 T-antigen. We observed no teratoma formation with C1EmMEn pre-iPSCs.

### 2.3 Comparative transcriptomics

Upon establishment of our iPSC lines and their differentiation capacity, we performed a comparative analysis of stem cells from a diverse set of other mammalian stem cells [10]. First, we looked at core pluripotency and naive pluripotency genes from humans, marmosets, mouse, rabbits, cattle, and elephants (**Figure 4a**). We found that while some of the expression of core pluripotency genes *OCT4, SOX2, NANOG*, and *LIN28A* was much lower in some of the emiPSC lines than all others, expression of some naive factors such as *TFAP2C, KLF4*, and *MYC* was higher than most. Furthermore, the general trend aligns well with the other large mammals like cattle and rhino. A principal component analysis of the overall transcriptome confirmed this finding (**Figure 4b**). Based on this natural clustering of species, we can group elephants into the zone of intermediate developmental clock according to analyses performed in the stem cell zoo study by Lazaro et al. [10].

**Figure 4:**
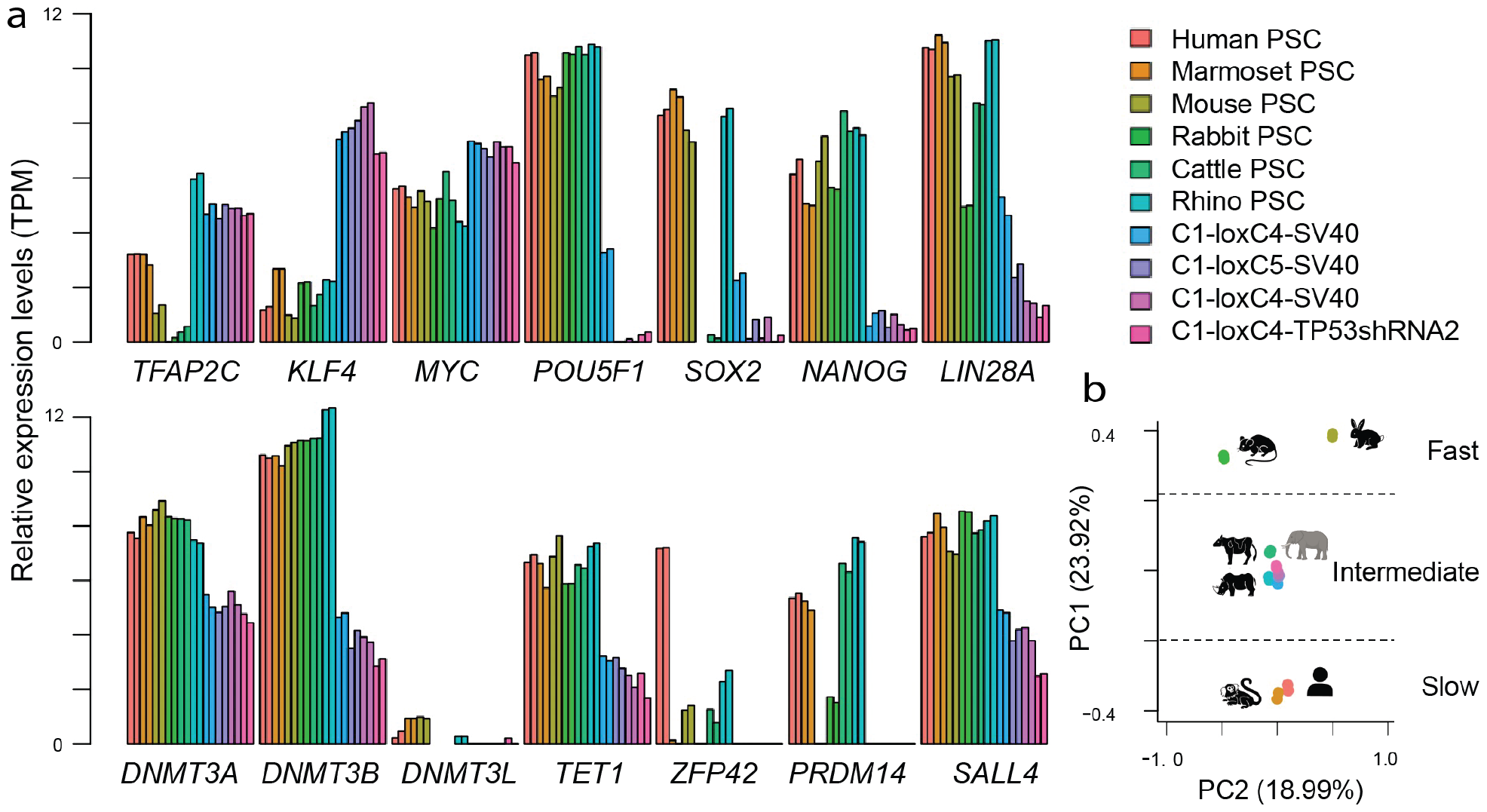
Distinct characteristics of elephant induced pluripotent stem cells. RNA-seq data of multiple species is compared. **a**. Comparison of canonical pluripotency gene expression in emiPSCs compared to iPSCs from other mammals. **b**. Principal component analysis of emiPSCs compared to iPSCs from other mammals. Data points clustered by approximate rate of developmental clock as described in [10].

## 3 Discussion

Here we demonstrated the first documented successful protocol for generating induced pluripotent stem cells in elephants. Our protocol required complex chemical media formulations to partially reprogram primary cells, and elephant-specific core pluripotency factors and modulation of cell growth pathways fully reprogram. Furthermore, we validated emiPSC karyotypic integrity, measured their high growth rate at negligible spontaneous differentiation rates, and determined that these cells have significant upregulation of many naïve pluripotency markers and some canonical primed markers.

We then functionally characterized these emiPSC lines and showed that they were able to form EBs and differentiated into the three germ layers. Additional EB immunostaining efforts are ongoing to resolve through EB 3D structures (i.e. 2D-imaging slices) and with additional markers. IF for tri-lineage differentiated cells, existing imaging, RNAseq, and RT-qPCR data all affirms that early differentiation markers are present in these experiments. Next, we showed that these lines are also capable of producing teratomas with diverse cell type-populations within, indicating differentiation potential. Since teratoma growth dynamics in vivo vary (even as long as 12 weeks), we are further validating the differentiation potential from an independent injection and longer-term monitoring followed by harvesting for analysis. Interestingly, naked mole rats have cancer resistance features and some cell cycle regulation irregularities shared with elephants (up-regulation of *CDKN1A/B* and others) that required modulation for the formation of teratomas [11]. We highlight these similarities for further studies into tumor formation for the purposes of pluripotency assessment and beyond.

This study provides the first successful report of producing emiPSCs, and it marks the beginning of more follow-up work. Further analysis into the implication of the *TP53* pathway in road-blocking elephant pluripotency is required. Of particular interest is disentangling the canonical versus the retrogene *TP53* contribution to determine the molecular features conferring reprogramming resistance.

The duration of reprogramming, which tends to be 5-10 days for model organisms like mouse, and over 3 weeks for large mammals including humans, was longer for the Asian elephant, at 2 months. We are further exploring whether alternative strategies can mitigate the longer reprogramming process observed in this study. For instance, in the study deriving naked mole rat iPSCs [39], the authors reduced reprogramming time by almost half, from one month to just slightly over two weeks.

Comparative analysis across a diverse set of mammalian iPSCs showed that our cells seem to cluster closest to other large-massed bodied mammals. However, observed differences in pluripotency marker expression and differentiation potential between species merit a deeper look into whether reprogramming methods or species differences are responsible for these observations. For example, the initially relatively low expression of *NANOG* was puzzling and boosting elephant iPSCs with more OSKM and *NANOG* did increase the overall endogenous *NANOG* and *SOX2* expression. We are further exploring whether this boost translates to more functional observations in ‘stemness’ and differentiation potential. Additionally, as our transcription-factor reprogramming constructs are excisable by design, we are validating the long-term effect on emiPSC endogenous pluripotency network following DOX withdrawal and transgene excision.

For conservation purposes and gametogenesis studies, validating truly naïve iPSCs is a critical step. Exploring the features of elephant stem cells that are unique to elephants, and endow cell-derived embryo models of development *ex utero* in a similar fashion to pioneering work in mouse and human embryos [40, 41], can shed much needed light in the gestation of complex higher mammals.

Bio-banking requirements such as freshly derived cells and choice of the parental cell lineage can also predict successful reprogramming outcomes, which further build the foundation of the conservation pipeline. Following this work, additional efforts into generating iPSCs from all extant species and subspecies of Proboscideans is critical for their conservation and understanding the similarities and differences between these iconic megaherbivores.

## 4 Methods

### RNA sequencing

RNA extraction was performed using the RNeasy Plus Universal Mini Kit (Qiagen 73404). After RNA preparation, RNA was sent directly to NovoGene for external QC and sample sequencing. For ATAC-seq sample, a frozen cell pellet was sent to NovoGene and all assays, library prep, and sequencing was performed by NovoGene.

### NGS analysis

RNA-seq analysis was performed using the Expression Analysis in RNASeq workflow on the Form Bio platform. Reads are trimmed using TrimGalore, to remove low quality (qual < 25) ends of reads and remove reads < 35bp. Trimmed reads are aligned to a reference genome using STAR2 (default) or HiSAT. BAMs from the same sample generated by multiple runs are merged using Samtools. The abundance of transcripts and genes are assessed using FeatureCount to generate raw gene counts, StringTie to generate FPKM and Salmon to generate raw transcript counts. Sample comparisons and differential gene/transcript expression analysis are performed using EdgeR, DESeq2 and IsoformSwitchAnalyzeR. ATAC-seq analysis is performed in a similar workflow. Specifically, reads are trimmed using TrimGalore, to remove low-quality (qual < 25) ends of reads and remove reads < 35bp. This workflow can be run with native open-source tools (NOST) or with Parabricks. With NOST, trimmed reads are aligned to a reference genome using BWA mem or Minimap2. BAMs from the same sample generated by multiple runs are merged using Samtools. Alignment quality is assessed using FastQC, Samtools, and Bedtools. With Parabricks, trimmed reads are aligned, duplicate reads are marked and alignment quality is accessed using fq2bam. Quality metrics are summarized with MultiQC. MA plots generated with DESeq2 (v3.18) and ggpubr (v0.6.0).

### Comparative transcriptomics

Public RNA-seq data for human, marmoset, mouse, rabbit, cattle and rhinoceros [10] were downloaded from NCBI Short Read Archive (SRA). We used GRCh38, mCalJa1.2.pat.X, GRCm38, OryCun2, BosTau9, NRM-Dsumatrensis-v1 as human, marmoset, mouse, rabbit, cattle, and rhinoceros reference genome, respectively. The count matrices from all species were subsequently merged by homologous gene sets that were downloaded from BioMart. Technical biases across datasets were then minimized by RUVg function with top 5,000 empirical controls in RUVSeq Bioconductor package (v1.28.0)

### Sendai virus reprogramming

*Elephas maximus* endothelial and epithelial cells were transduced with Sendai virus (Thermofisher Scientific, A16517) following manufacturer’s instructions, with an MOI of 5:5:3 and an MOI of 10:10:6. Different cell densities were tested, with 1 × 10^5^, 1.5 × 10^5^ and 3 × 10^5^ cells transduced per reaction, with the addition of 5*μ*g/mL of protamine sulfate (Sigma, P3369-10G). Cells were seeded on 3T3-J2 cells (StemCell Technologies: 100-0353) and Geltrex (Thermofisher Scientific, A1413302).

### Lentivirus reprogramming

*Elephas maximus* endothelial and epithelial cells were transduced with Lentivirus (Sigma, SCR5451) following manufacturer’s instructions, with an MOI of 2, 5, 10, and 20. A total of 1 × 10^5^ and 1.5 × 10^5^ cells were transduced per reaction, with the addition of 5*μ*g/mL of protamine sulfate (Sigma, P3369-10G). Cells were seeded on 3T3-J2 cells (StemCell Technologies: 100-0353) and Geltrex (Thermofisher Scientific, A1413302). Lentivirus reprogramming (Sigma, SCR5451) with an MOI of 10 was also tested in Elephas maximus endothelial and epithelial cells, following manufacturer’s instructions, with the addition of transcription factors *OCT4, LIN28A*, and *NANOG* (Cellomics, PLV-10012-50, PLV-10015-50, and PLV-10075-50 respectively), each at an MOI of 10. A total of 1 × 10^5^ and 1.5 × 10^5^ cells were transduced per reaction, with the addition of 5*μ*g/mL of protamine sulfate (Sigma, P3369-10G). Cells were seeded on 3T3-J2 cells (StemCell Technologies: 100-0353) and Geltrex (Thermofisher Scientific, A1413302).

### Cellular reprogramming

Primary *Elephas maximus* endothelial cells were used as the starting cell line for reprogramming. These cells are mantained in 30% FBS/1% antibiotic/antimycotic /1% Non-essential AA/ EGM-2 media (Lonza) with Laminin521 coating (5 *μ*g/ml; Gibco A29248). Once the cells became approximately 70% confluent, a chemical cocktail medium was used for partial reprogramming to emPRCs - KO DMEM (Gibco 10829-018) + 10% KOSR (Invitrogen) + 55 *μ*M 2-mercaptoethanol (55 mM (1000X); Gibco 21985023) + 50 ng/ml bFGF (20 *μ*g/ml; heat stable; Life technologies PHG0369) + 0.5 mM VPA (EtOH; Selleckchem S3944) + 5 *μ*M CHIR-99021 (DMSO; Selleckchem S1263) + 2 *μ*M RepSox (DMSO; Selleckchem S7223) + 10 *μ*M Tranylcypromine (2-PCPA) HCl (DMSO; Selleckchem S4246) + 20 *μ*M Forskolin (DMSO; Selleckchem S2449) + 1 *μ*M Ch 55 (DMSO; Tocris 2020) + 5 *μ*M EPZ004777 (DMSO; Selleckchem S7353). The medium was changed every two days until small emPRC colonies were observed. Once colonies reached sufficient size, they were hand-picked and mechanically passaged 2-3 times. Once a 2-3 million emPRCs were avialable, they were necleofected with plasmids encoding genome-integrating (via Piggy-Bac), inducible, polycistronic transgene expression cassettes. These cassettes contained one of [*Loxodonta africana OCT4/SOX2/KLF4/CMYC* (C4) or *Loxodonta africana OCT4/SOX2/KLF4/CMYC/LIN28A* (C5) or *Loxodonta africana OCT4/SOX2/KLF4/CMYC/LIN28A/NANOG* (C6)] and [SV40 T-antigen or an shRNA targeting *TP53* retrogenes in *Elephas maximus*]. Cells were recovered, selected with mammalian selection markers hygromycin and puromycin, and induced for 1 month until full emiPSC morphology, growth, and molecular signature was observed.

### Immunofluorescent staining

EBs were fixed using 4% v/v paraformaldehyde (15710, Electron Microscopy Sciences, Hatfield, PA, USA) for 20m, washed three times and permeabilized with 0.5% Triton X-100 (Sigma-Aldrich, Saint Louis, MO), for 20 min. Samples were washed three times and blocked with 5% BSA, 0.05% Triton for 1 h and incubated with primary antibodies diluted in 1% BSA, 0.05% Triton-X100 overnight at 4C. Samples were washed three times and incubated with secondary antibodies for 1h at 37C, then washed three times and counter-stained with Hoechst 33342 (H1399, Invitrogen, Carlsbad, CA, USA) for 10m at RT, washed three times and mounted with Vectashield (Vector Labs., H100010A). Micrographs were acquired with a Zeiss AxioObserver .5 LED fluorescent microscope, High Performance microscopy camera Axiocam 705 mono R2. Objective LD A-Plan 10X/0.25 Ph1. emiPSCs were stained with custom elephant-specific antibodies for *NANOG* and *OCT4* with TXR-anti-rabbit (Invitrogen T-2767) and *SOX2* (Invitrogen MA1-014) with AF488-anti-mouse (Invitrogen A-21202). EBs were stained with for *GATA4* with a custom elephant-specific antibody, *PAX6* (Invitrogen MA1-109), and *FOXA2* (Novus NB100-1263). Secondary antibodies used were TXR-anti-rabbit (Invitrogen T-2767), TXR-anti-goat (Invitrogen PA1-28662), and AF488-anti-mouse (Invitrogen A-21202).

### Embryoid body formation

Embryoid bodies (EBs) were formed using either AggreWell 400 plates (StemCell 34450) or ultra low attachment 96 well plates (Corning 4515). Plates were pre-treated with Anti-Adherence Rinsing Solution, after which approximately 400 cells/microwell for AggreWell 400 or (5k/well) for 96-well plates in the described reprogramming medium above were added and centrifuged at 100g for 3m. Plated cells were left undisturbed for 24h at 37C with 5% CO2 and 95% humidity. After 24h, medium was changed every 48h with either AggreWell EB Formation Medium (StemCell 05893) or reprogramming medium without disturbing the cells, prior to IF and imaging.

### Tri-lineage differentaition

50,000 emiPSCs were plated onto a Laminin521 coated 12 well plates and treated for 10 days with the STEMdiff Trilineage Differentiation Kit (StemCell 05230).

### Chromosomal Isolation and Counting

Chromosomal isolation and counting was performed by incubating cells in for 3h at 37C. Afterwards, cells were resuspending cells in 0.075M KCl solution at 37C for 8m. Next, cells were resuspended in 1ml of fixative and gently mixed and incubated at RT for 10m. Cells were then centrifuged at 900rpm for 8m and resuspended in a fixative at RT for 10m. This fixation step was repeated twice. Finally cells were mounted onto a slide with dye for imaging.

### Teratoma

200,000 - 1,000,000 emiPSCs were injected into the hindleg of immuno-compromised mice. Legs were observed daily for 6 weeks, after which tumor masses were extracted. Teratoma sections were evaluated by professional histologists at Histowiz.

## 5 Acknowledgements

The authors would like to acknowledge Beth Shapiro for helpful comments on the manuscript, and thank the members of the Colossal Biosciences Scientific Advisory Board, Conservation Advisory Board, Executive Advisory Board, and Executive Core Leadership for their conversations and support on this effort. We would also like to thank Colossal’s global academic and industrial partners. We thank BioRender for visualization and AbClonal for custom antibody design.

## 6 Author Contributions

Conceptualization, E.H.; methodology, E.H., and E.A.; formal analysis E.A., K.H., C.R.C., Y.T., K.B., C.R.I., N.K., H.B., X.A.P., J.N.s A.K., B.C. and E.H.; investigation, E.A., K.H., C.R.C., Y.T., A.A.T., K.B., C.R.I., N.K., H.B., X.A.P., A.Q., G.R., G.K., M.M., N.M., K.Z., J.K., A.B., B.M., and E.H.; resources B.L. and G.C.; data curation, E.A., Y.T., K.B., and C.R.I.; writing, E.A. and E.H.; visualization, E.A. and E.H.; supervision, E.A., B.C., B.L., G.C., and E.H.; project administration, E.A. and E.H.; funding acquisition, B.L. and G.C.

