## Supplementary Information for "Derivation of elephant induced pluripotent stem cells"

### Derivation of elephant induced pluripotent stem cells Supplementary Information

#### 1 Partial emiPSC Reprogramming Attempts

The process of creating emiPSCs required the testing of many established reprogramming methods that have been tried and tested on other species. While many of these attempts resulted in no morphologial change to the starting cell population, cell population senescence, and cell death, some protocol were able to produce cells that appeared partially on their way to iPSC characteristics (**Supplementary Figures 1-9**). Briefly, we attempted to (1) screen broad new sets of human TF combinations with the human TFome (**Supplementary Figure 1**), (2) create emiPSCs via standard viral reprogramming methods utilizing human transgenes (**Supplementary Figure 2, 3**), (3) induce cellular reprogramming via complex chemical treatment progressions supplemented with episomal transgene expression [1] on two cell lines (one male, one female) (**Supplementary Figure 4a,b**), (4) create emiPSCs from polycistronic PiggyBac mouse OSKML and SV40T antigen vectors (**Supplementary Figure 4c**), (5) create emiPSCs from polycistronic PiggyBac elephant OSKM/NL and SV40T antigen or TP53 shRNA vectors (**Supplementary Figure 4d**), and (6) create emiPSCs via chemical induction alone (**Supplementary Figure 4e**).

One of our first attempts to reprogram these notoriously difficult cells was to question whether or not we were missing additional reprogramming TFs that may not yet be described in the literature. To this end, we applied computational tools to NGS data from human iPSCs and ESCs [2, 3, 4] (**Supplementary Table 1**). We tested these combinations with these sets of TFs and enriched populations of cells that exhibited potential up-regulation of cell surface pluripotency markers (**Supplementary Figure 1a**). While we found some interesting enrichment in these samples (**Supplementary Figure 1c-e**), testing of these refined combinations did not yield further success - cells in these protocols tended to form small, tight colonies at the beginning of the experiment and then these colonies stagnated (**Supplementary Figure 1b**). We speculated at this point whether or not elephant TF amino acid sequences are different enough that human TF proteins may not work as they are supposed to in these cells, but as most TF-binding domains of TFs tend to be conserved, we still have no conclusive data on this topic. It was also of concern that since all of these TFs were on individual expression cassettes, that a screen may miss requirements where all members of a reprogramming set are required.

Accordingly, we hypothesized that polycistronic format may be required. As a first test of these class of methods, we applied standard polycistronic and monocistronic viral reprogramming methods (**Supplementary Figure 2**). While these methods showed some clear shifts in cell morphology at relatively quick time scales, the attempts again resulted generally in a combination of cell senescence and cell death (**Supplementary Figure 3**).

Next, we attempted to reprogram emECs with polycistronic PiggyBac expression Yamanaka factors (**Supplementary Figure 4c**). The female endothelial cell line we tried to reprogram solely with trans-gene over-expression via PiggyBac yielded a different morphology than the other two (**Supplementary Figure 6c**), but alas still did not show high pluripotency marker expression, aside from *SOX2* (**Supplementary Figure 7b**). While the morphology of these colonies was loosely on track, the populations grew very slow and still lacked key characteristics of iPSCs. In parallel, we also tested a similar method that used elephant-specific OSKM/NL amino acid sequences and shRNAs to modulate the expression of *TP53* and its retrogenes (**Supplementary Figure 4d, 5**). While we could validate that the shRNAs were repressing either the canonical *TP53* sequence and its retrogenes (*TP53shRNA4*) or just most retrogenes (*TP53shRNA2*) (**Supplementary Figure 5**), this transgene-only approach was still not effective at creating emiPSCs (**Supplementary Figure 6d, 7d**).

Following this set of attempts, we ventured into testing methods that combined both transgene and complex media formulations (**Supplementary Figure 4a,b**). These attempts used complex chemical treatments, supplemented with episomal reprogramming factors over-expression. After similar treatments, the female line showed marker profiles and morphology roughly characteristic of cells in the trophoblast stem-cell-like state (**Supplementary Figure 6b, 7c**), and the male line showed marker profiles and morphology roughly characteristic of cells undergoing the mesenchymal to epithelial transition (MET) (**Supplementary Figure 6a, 7a**).

Finally, attempts to reprogram elephant cells with the same chemical protocol demonstrated for mice [5] were interesting in both a morphological (**Figure 1e-g**) and growth perspective. We tested a number of different variations of chemical cocktails - A (0.5 mM VPA, 10uM CHIR, 20uM Repsox, 10uM Tranly, 10uM Forskolin, 1 RAR agonist), B (0.5 mM VPA, 15uM CHIR, 2uM Repsox, 10uM Tranly, 20uM Forskolin, 2 RAR agonist) and C (0.5 mM VPA, 10uM CHIR, 10uM Repsox, 5uM Tranly, 10uM Forskolin). While these features were exciting, molecular analysis revealed that these cells still did not have expression of core pluripotency factors (**Supplementary Figure 8**). While this result was puzzling, given the morphology, growth and expression of secondary pluripotency factors, we pursued adding PiggyBac, polycistonic Yamanaka factors using the endogenous *Loxodonta africana* sequences and modulators of TP53 and/or its retrogenes to these cells (**Supplementary Figure 4f**). Here we note that chemical cocktail B was the chosen cocktail to pursue a majority of TF-overexpression experiments. While the NGS data suggested that chemical cocktail C (**Supplementary Figure 9**) was indeed closer to final emiPSC molecular composition, it grew slower that cells derived from cocktail B, and that with a repeated process with refined colony-picking strategies, chemical cocktail B ultimately produced the cells we call throughout the manuscript 'C1EmMEn pre-iPSC'.

Principal component analysis of all partial success cell lines and final success cell lines paints a very interesting picture (**Supplementary Figure 9**). Surprisingly, the pure polycistronic transgene expression method that utilized SV40T as opposed to TP53shRNAs clustered very closely to the hybrid chemical/transgene methods and indeed not far of from true emiPSCs. Similarly, purely chemical methods appeared to do a lot of heavy lifting to make emiPSCs, but could not alone finish the job. Even more striking was the case of C3-loxc4-TP53shRNA4 cells, which appeared to create some morphological changes and had many of the needed components to create emiPSCs, yet appears to have barely moved the needle towards emiPSCs and apparently in the wrong direction.

#### 2 Additional emiPSC Characterization

Our emiPSC lines share a vast majority of molecular, morphological, and phenotypic features, however there are some minor differences. These are exemplified with small variations to teratoma size and composition (**Supplementary Figure 16**), some features related to growth (**Supplementary Figure 17**), and embryoid body features (**Supplementary Figure 18**). We also show that expression of early differentiation markers increases over time during EB formation, as expected (**Supplementary Figure 14**). All primers for all qPCRs shown in (**Supplementary Table 2**).

Intriguingly, while most iPSCs from other well-studied organisms exhibit a strong down-regulation of *TP53*, *CDKN1A*, *CDKN1B*, *CDKN2A*, and *CDKN2B*, emiPSCs appear to to do the opposite - in most cases these genes are either up-regulated or unchanged compared to primary cells. We intend to explore this fully in future work to determine why elephant cells are so difficult to reprogram.

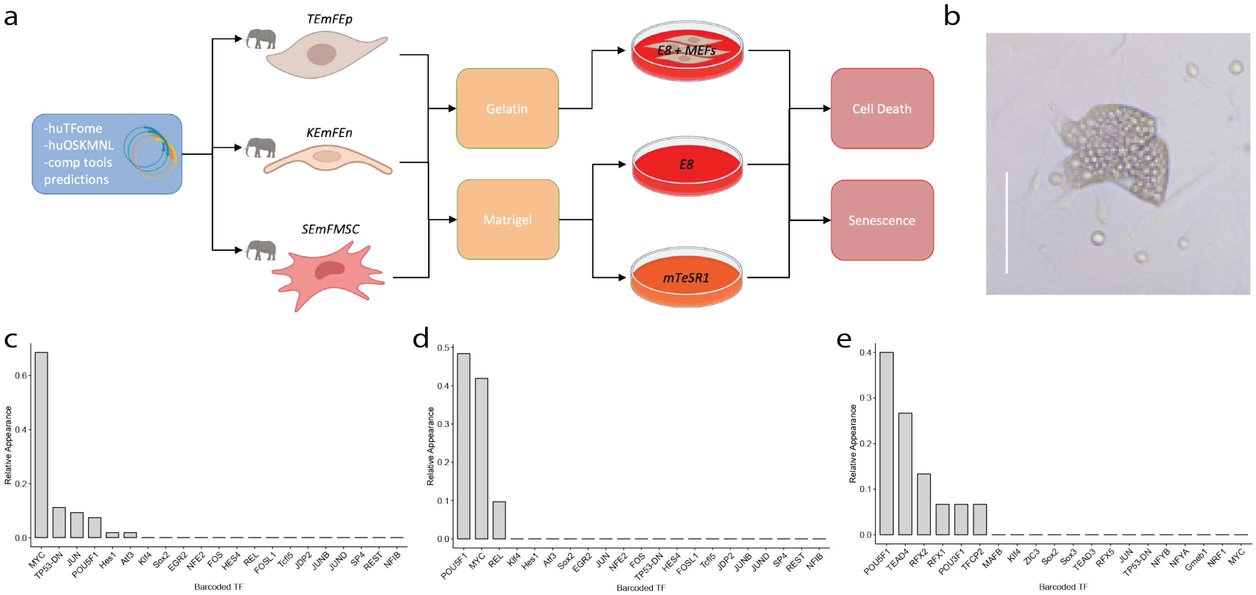


**Supplementary Figure 1: TFome TF-enrichment screening. a.** TFome screening was performed on emECs and emMSC (*Elephas maximus* mesenchymal stromal cells) on a small set of simple reprogramming conditions. These attempts all resulted in either cell death or **b.** heterogenous appearance of senescent colonies. These cell populations were FACS-sorted and sequenced to determine enrichment of TFs used to produce these cells. All cells shown at 10X magnification, scale bar = 200$\mu$M. Relative TF enrichment is shown for a set of TFs identified from human iPSCs that stained for **c.** *SSEA1* and **d.** *TRA-1-60* and TFs identified from human ESCs for **e.** *SSEA1*. Subsequent rounds of screening with these refined sets of TFs did not yield improvement.

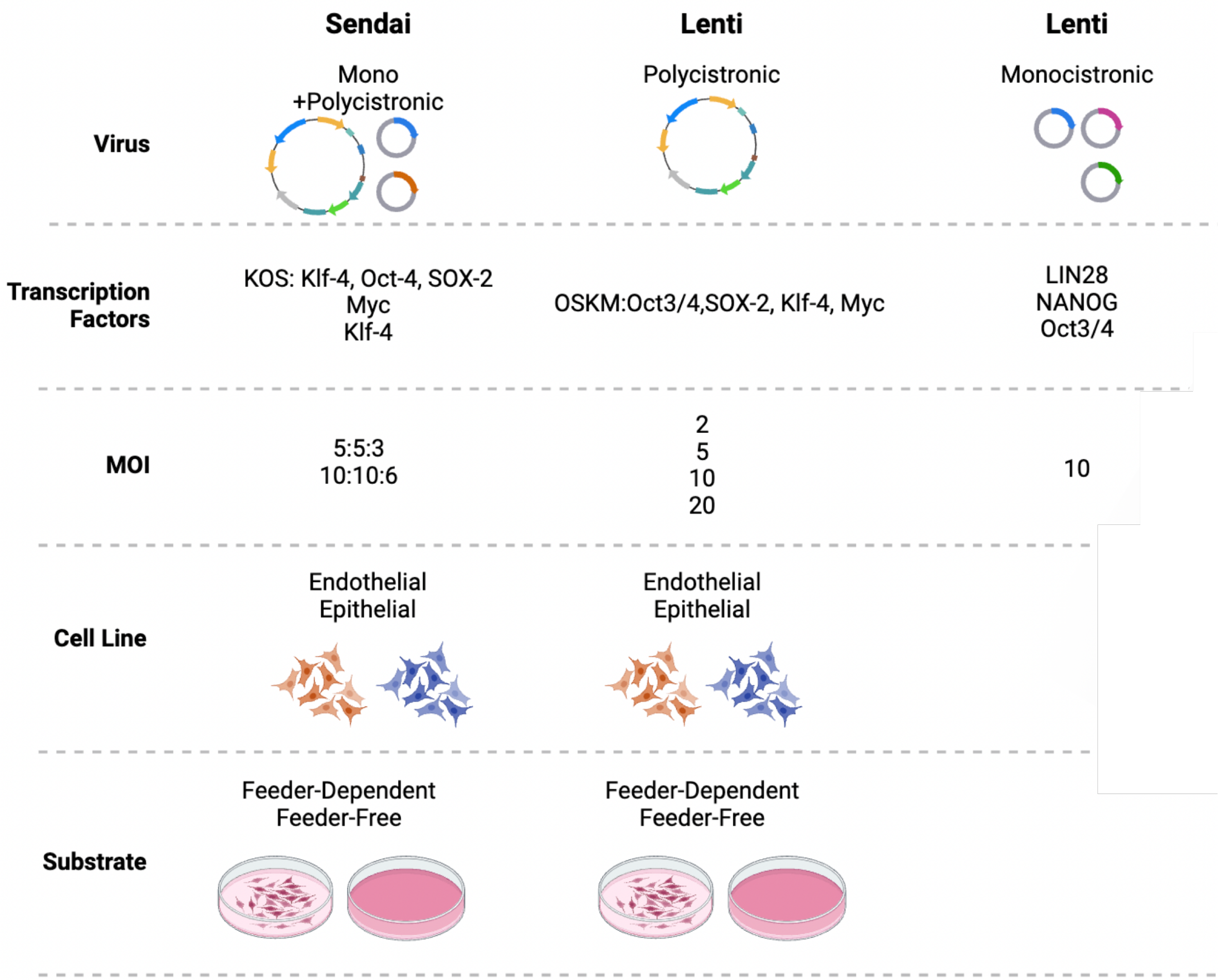


**Supplementary Figure 2: Viral reprogramming attempt conditions.** emECs were treated with viral transductions with both Sendai virus and Lenti virus under standard reprogramming conditions defined by these methods. Different MOIs and standard plating conditions were tested.

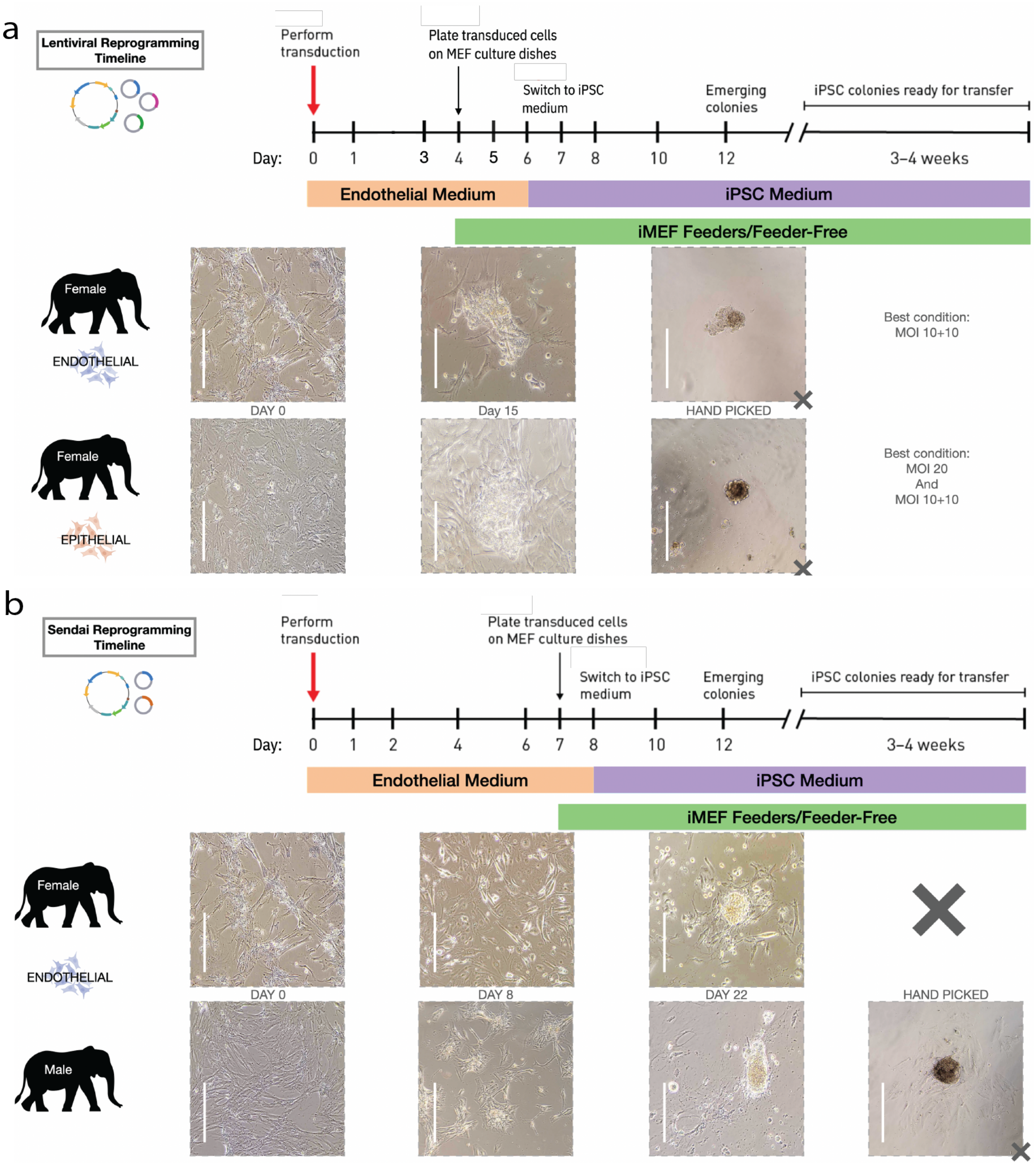


**Supplementary Figure 3: Viral reprogramming attempt morphological progression.** All cells shown at 10X magnification, scale bar = 200$\mu$M. **a.** Lenti-viral treatment reprogramming timeline and morphological results. **b.** Sendai virus treatment reprogramming timeline and morphological results. In both cases, senescent colonies emerged that died after attempted transfer.

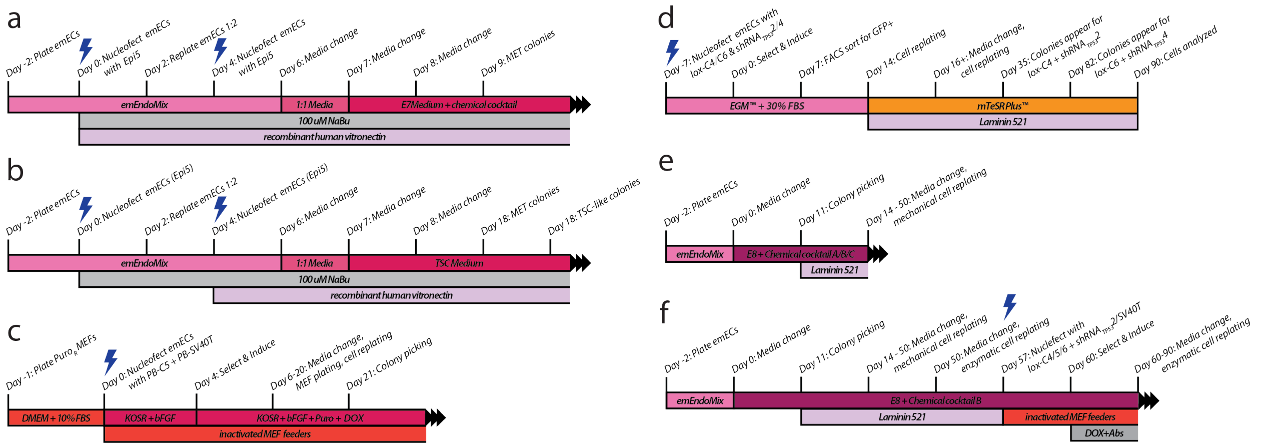


**Supplementary Figure 4: Culture conditions for TF-based, chemical, and hybrid reprogramming attempts.** A vast majority of reprogramming attempts resulted in no morphological change, cell death, or cell senescence. Some protocols yielded cells that were partially reprogrammed. Here we show the reprogramming protocols for: **a.** C2 MET cell line **b.** TEmFEp TSC cell line **c.** KEmFEn TF-based polycistronic reprogramming cell line **d.** C3-loxC4-TP53shRNA4 cells **e.** pre-iPSCs with either chemical cocktail A, B, or C; and finally **f.** the successful emiPSC protocol.

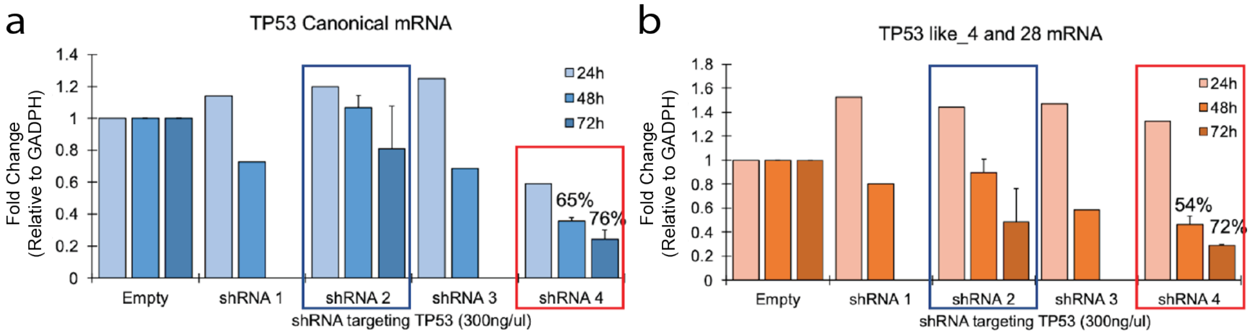


**Supplementary Figure 5: shRNAs targeting *Elephas maximus TP53* and its retrogene expansions.** A set of 4 shRNAs were designed to target TP53 and/or its 29 corresponding retrogenes (RTGs). We tested the shRNAs against **a.** the full length *TP53* mRNA sequence and **b.** 19 *TP53* RTGs including *TP53-RTG-4* and *TP53-RTG-28* mRNA sequences and measured fold change realative to an empty shRNA vector for 24, 48, and 72 hours.

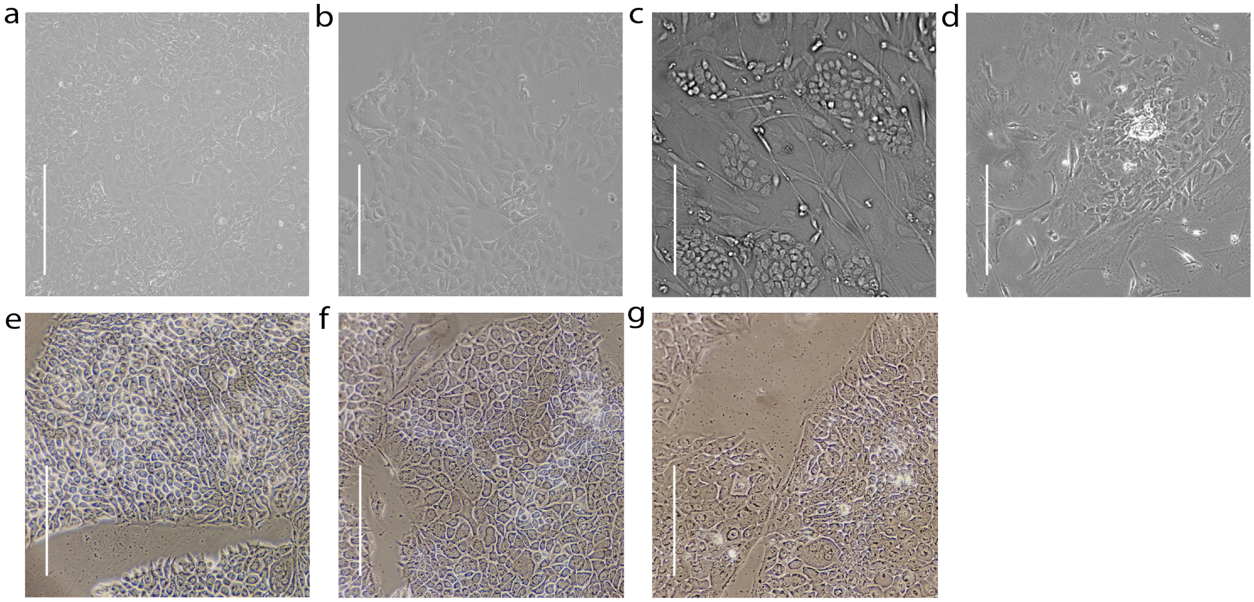


**Supplementary Figure 6: Cell morphologies for partial reprogramming.** Partially reprogrammed cells showed morphological features that were reminiscent of intermediate reprogramming morphologies. All cells shown at 10X magnification, scale bar = 200uM **a.** C2 MET cells. **b.** TEmFEp TSC cells. **c.** KEmFEn piPSC cells. **d.** C3-loxC4-TP53shRNA4 cells. **e.** Chemical cocktail A cells. **f.** Chemical cocktail B cells. **g.** Chemical cocktail C cells.

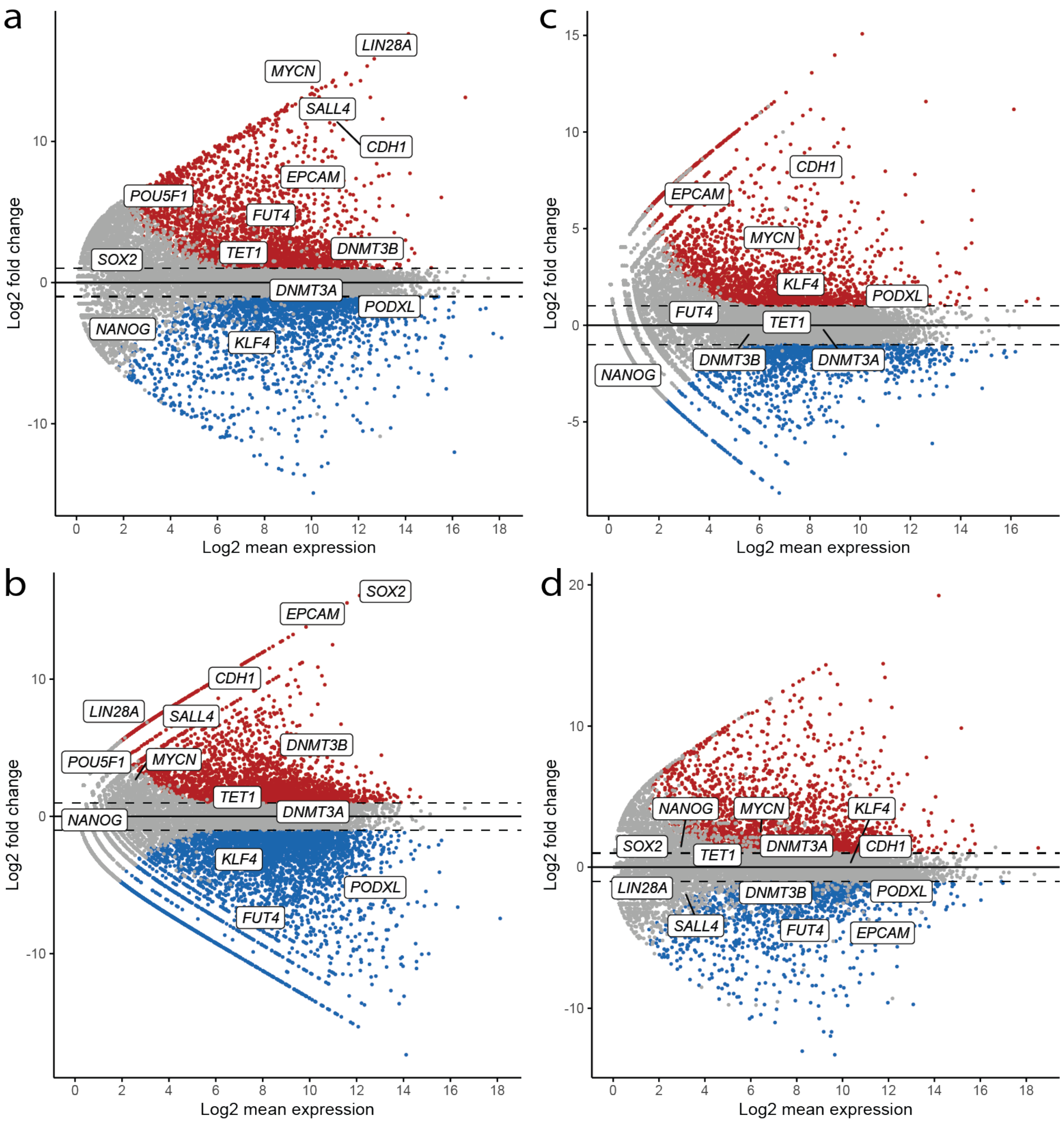


**Supplementary Figure 7: MA plots for partial reprogramming.** All plots show significantly up-regulated genes (red), significantly down-regulated genes (blue), and non-significant change (gray). **a.** C2 MET. **b.** KEmFEn piPSC. **c.** TEmFEp TSC. **d.** C3-loxc4-TP53shRNA4

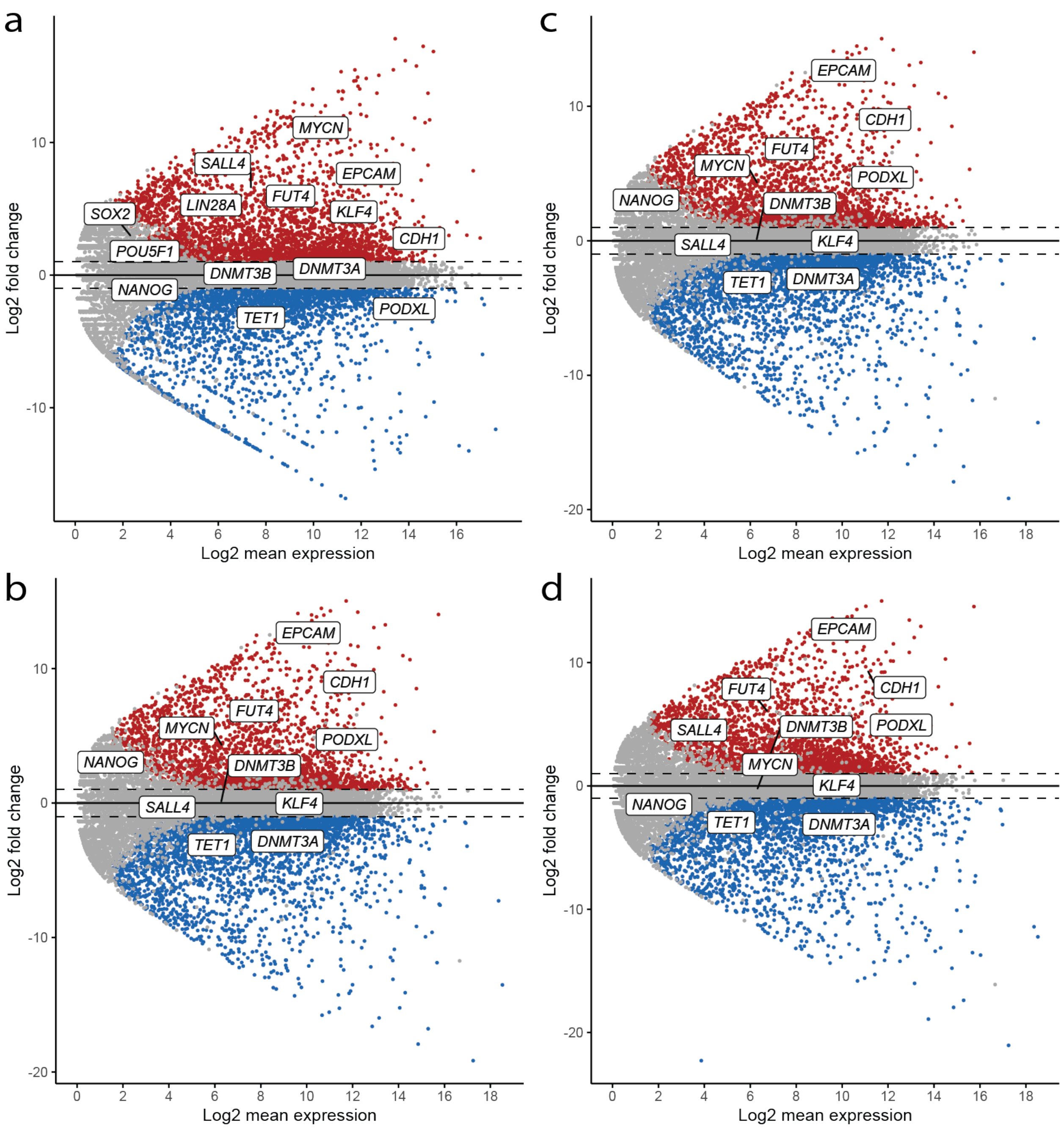


**Supplementary Figure 8: MA plots for partial reprogramming.** All plots show significantly up-regulated genes (red), significantly down-regulated genes (blue), and non-significant change (gray). **a.** C1EmMEn pre-iPSC **b.** C1EmMEn pre-iPSC A **c.** C1EmMEn pre-iPSC B **d.** C1EmMEn pre-iPSC C

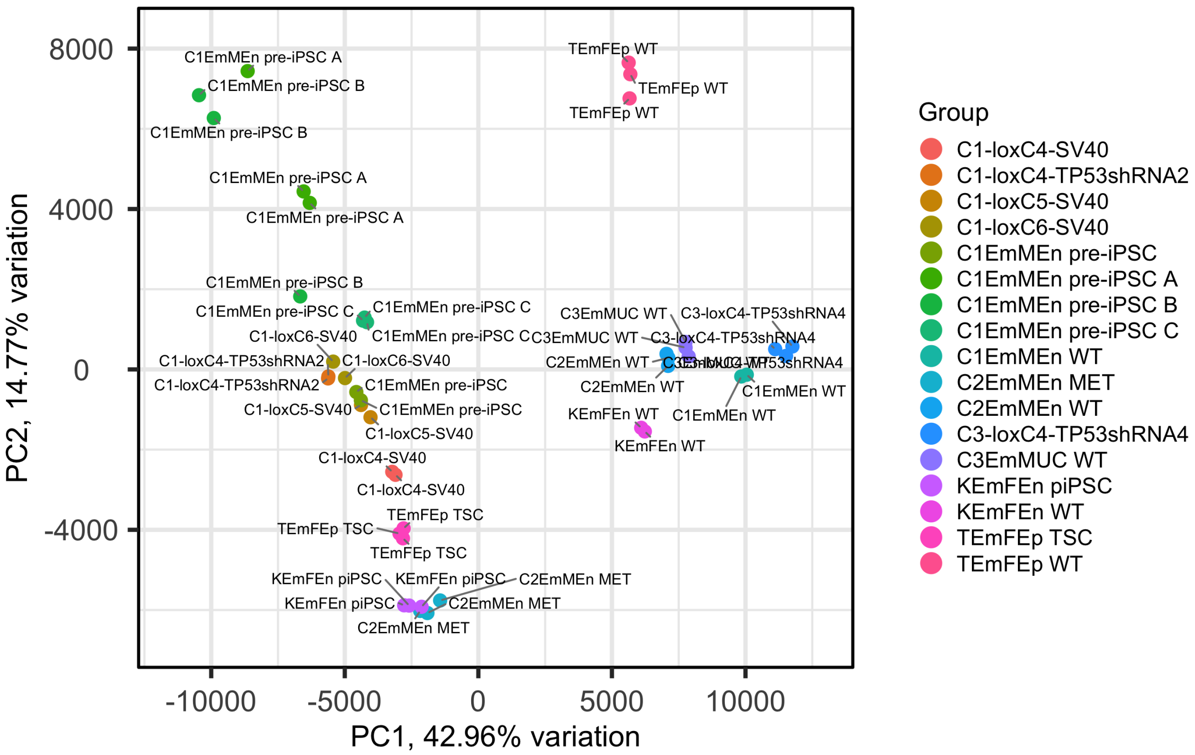


**Supplementary Figure 9: Principal component analysis of partial and full reprogramming.** PCA analysis compares all primary cell populations, partial reprogramming populations, and full reprogramming populations for the first two principal components (PC1, PC2).

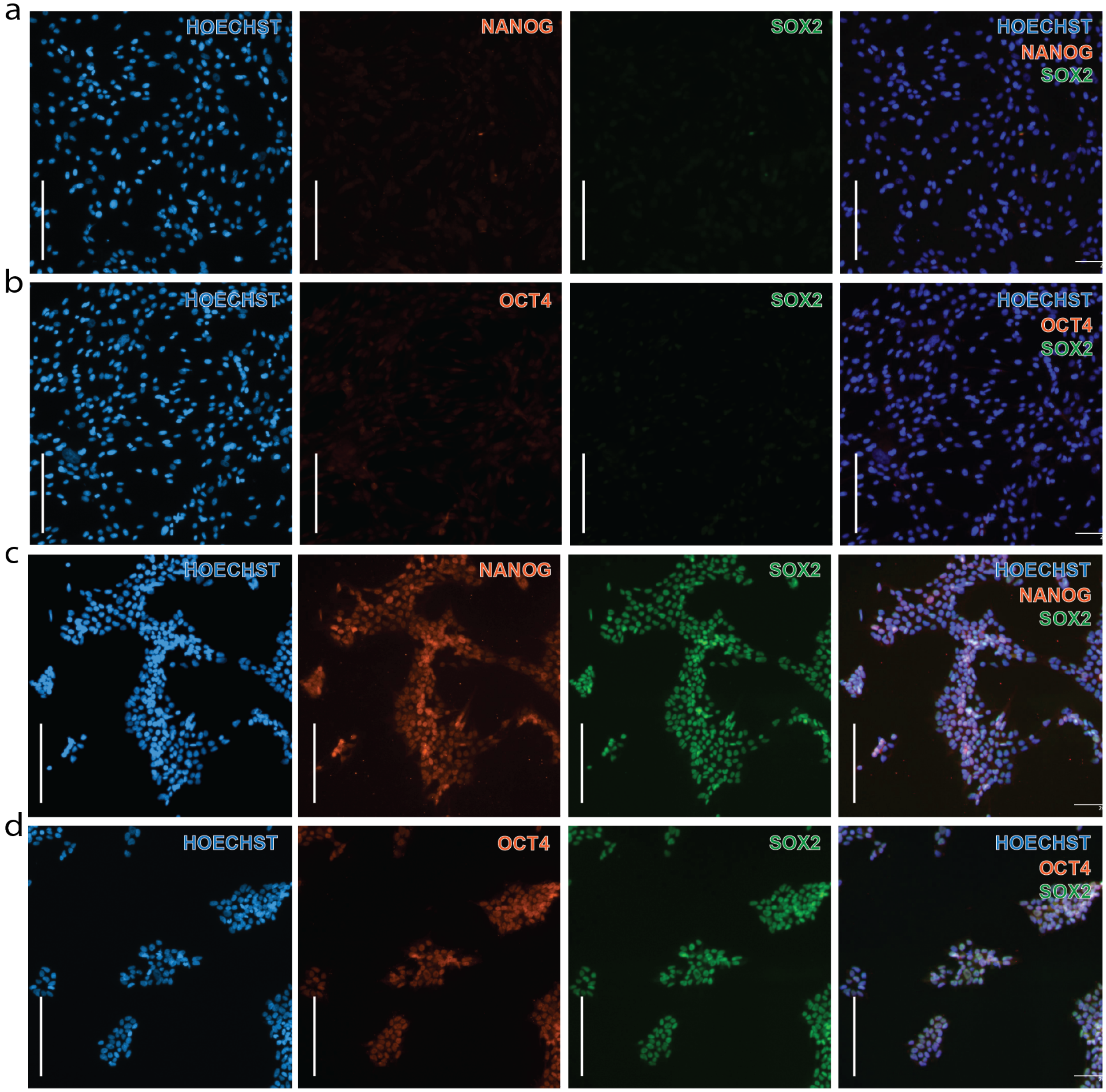


**Supplementary Figure 10: Immunofluorescent staining controls for pluripotency markers**. All scale bars = 200$\mu$M, magnification = 10X. *NANOG* and *OCT4* antibodies are custom elephant amino-acid sequence optimized, *SOX2* antibody is commercial due to extremely high sequence overlap. **a.** Immuno-fluorescent detection of *NANOG* (Texas Red) / *SOX2* (AF488) / HOECHST separately and merged in C1EmMEn WT cells. **b.** Immuno-fluorescent detection of *OCT4* (Texas Red) / *SOX2* (AF488) / HOECHST separately and merged in C1EmMEn WT cells. **c.** Immuno-fluorescent detection of *NANOG* (Texas Red) / *SOX2* (AF488) / HOECHST separately and merged in human iPSC cells. **d.** Immuno-fluorescent detection of *OCT4* (Texas Red) / *SOX2* (AF488) / HOECHST separately and merged in human iPSC cells.

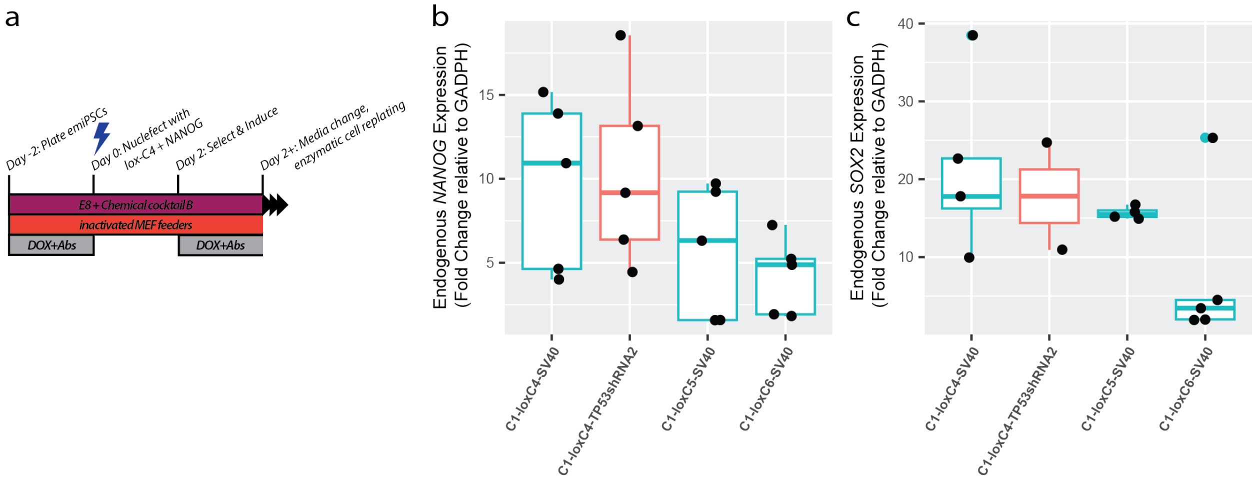


**Supplementary Figure 11: emiPSC lines reinforced with additional transgene expression. a.** emiPSC lines were reinforced with additional copies of loxC4 (OSKM) and *NANOG* PiggyBac overexpression cassettes via a serial transfection. Cells were then measured for relative expression changes after 7+ days via RT-qPCR for primer pairs flanking exon-intron regions of **b.** *NANOG* and **c.** the coding sequence and 3' UTR of *SOX2* to ensure that measured expression increases were endogenous and not simply transgene expression.

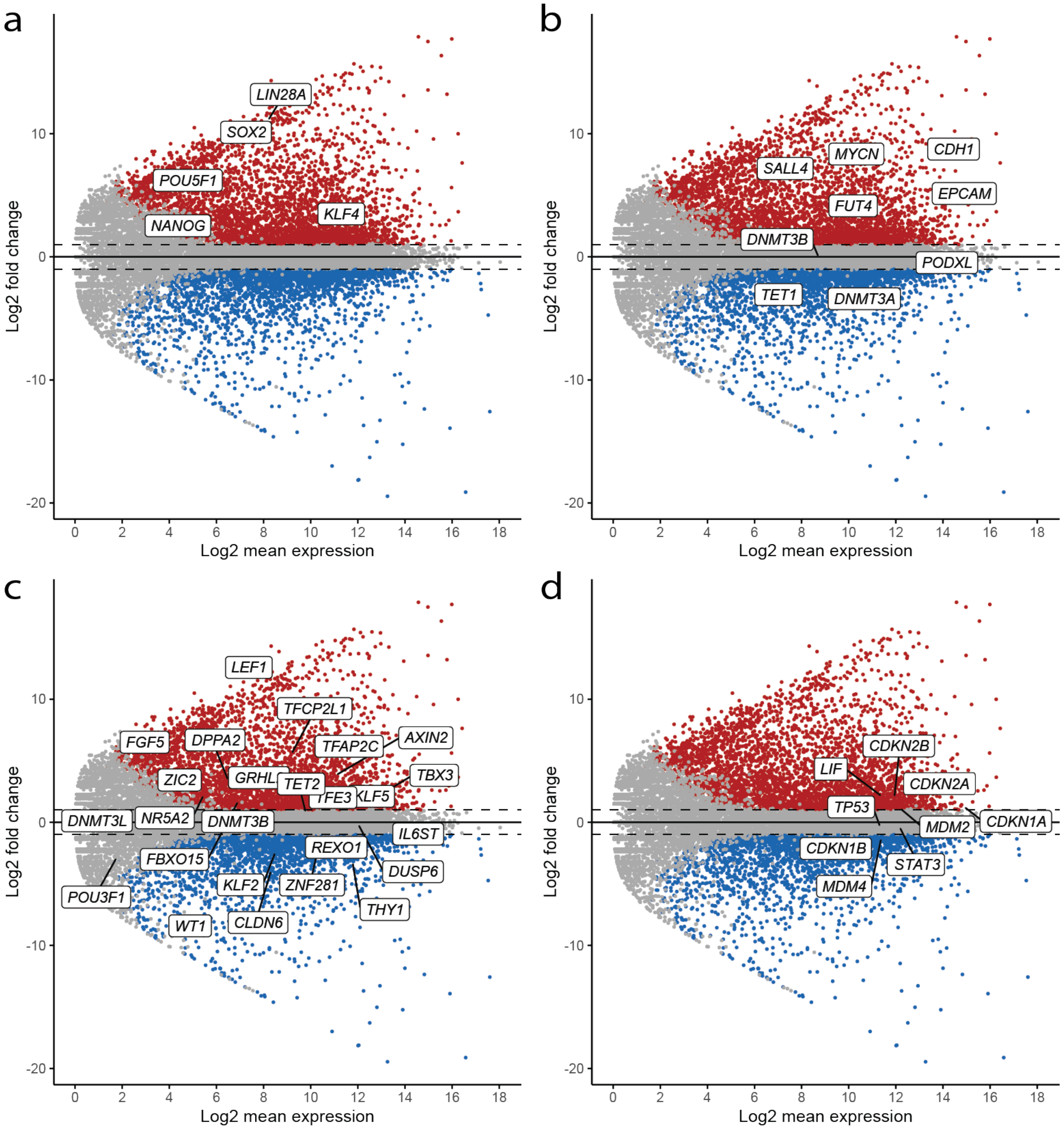


**Supplementary Figure 12: C1-loxC4-SV40 cell line markers after withdrawl of DOX for 10 days** MA plots shown for up- and down-regulation. All plots show significantly up-regulated genes (red), significantly down-regulated genes (blue), and non-significant change (gray). **a.** Core pluripotency markers **b.** Additional pluripotency markers **c.** Canonical naive and primed markers *d.* Core cell-cycle regulatory markers.

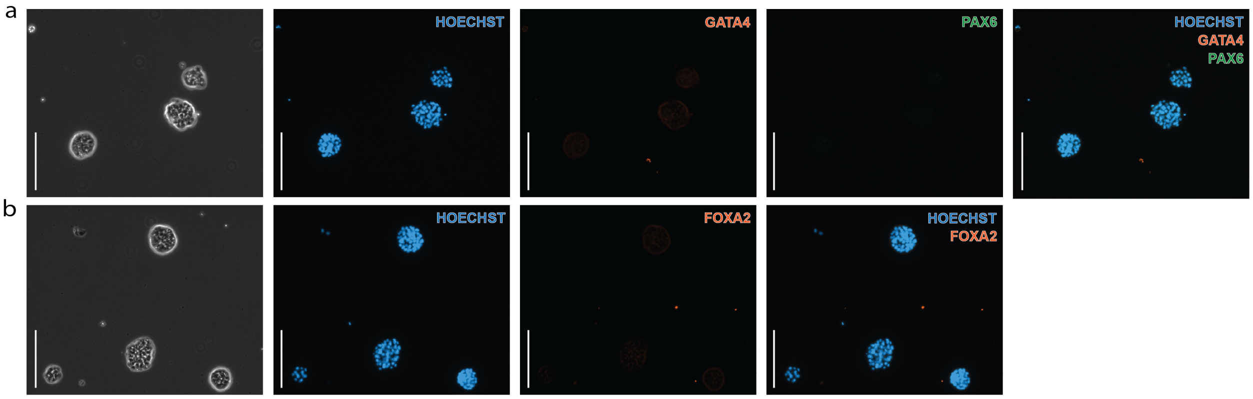


**Supplementary Figure 13: Immunofluorescent staining controls for early differentiation markers**. All scale bars = 200$\mu$M, magnification = 10X **a.** Immunofluorescence (IF) microscopy images of embryoid bodies (EBs) formed by emiPSC line C1-loxC5-SV40 stained with only secondary antibodies detecting the expression of lineage-specific markers *PAX6* (ectoderm) and *GATA4* (mesoderm). **b.** IF microscopy images of EBs formed by emiPSC line C1-loxC5-SV40 stained with only secondary antibodies for the expression of lineage-specific marker *FOXA2* (endoderm).

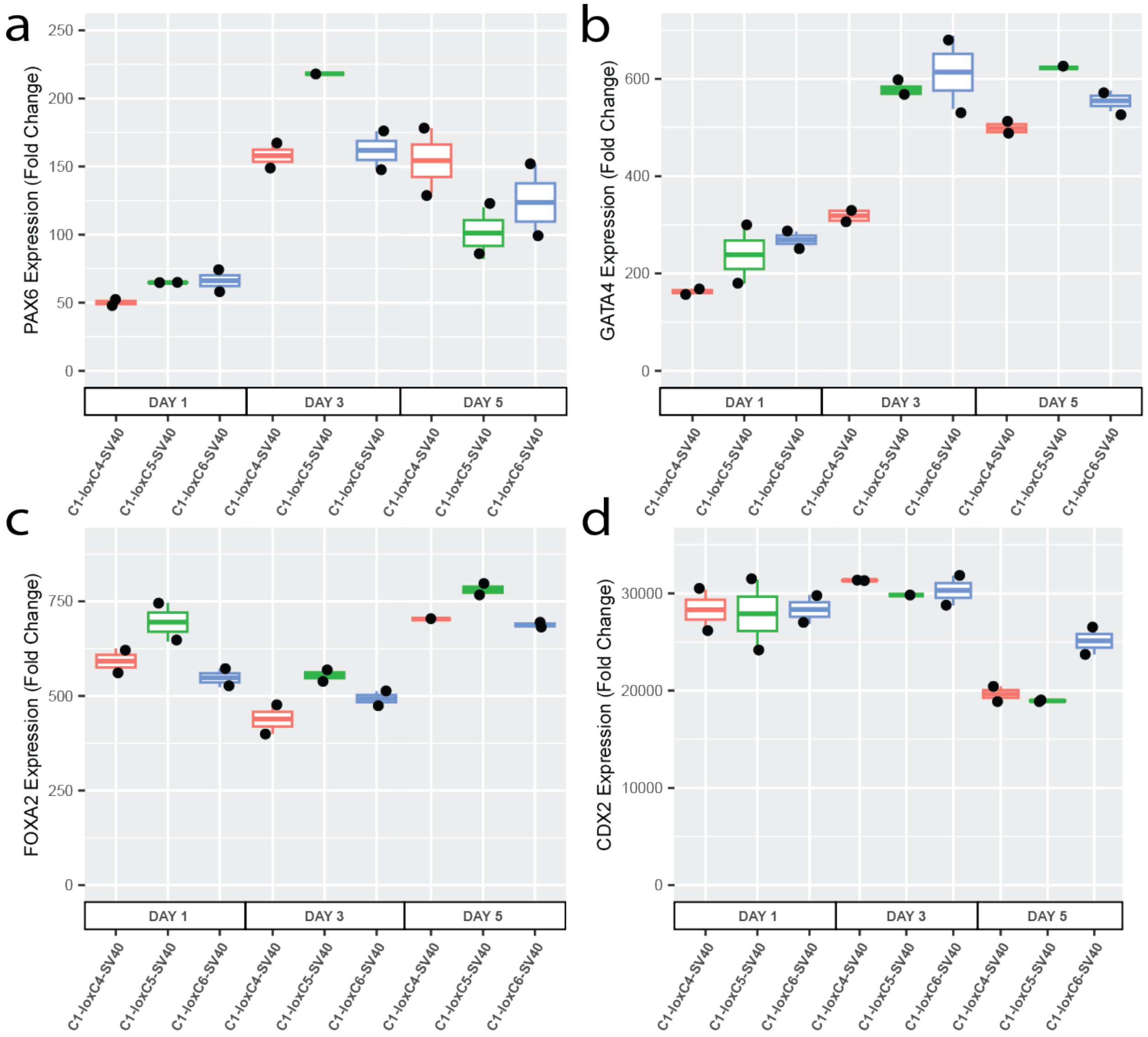


**Supplementary Figure 14: Upregulation of early differentiation markers in EBs over time.** Expression of early differentiation markers were measured over the course of EB formation for 3 emiPSC cell lines. Data shown for RT-qPCR fold change compared to C1EmMEn WT cells relative to GADPH expression. **a.** *PAX6* (Ectoderm) **b.** *GATA4* (Mesoderm) **c.** *FOXA2* (Endoderm) **d.** *CDX2* (Endoderm)

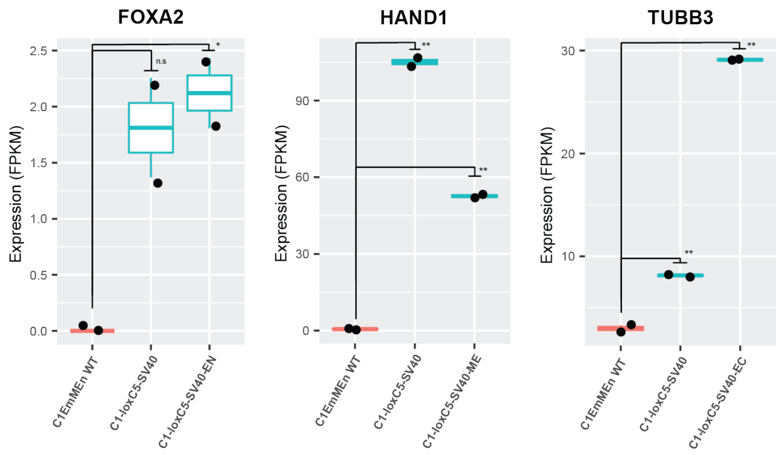


**Supplementary Figure 15: Additional tri-lineage differentiation markers analyzed via RNAseq**. Expression of early differentiation markers *TUBB3* (ectoderm), *HAND1* (mesoderm), and *FOXA2* (endoderm) following tri-lineage differentiation in emiPSC line C1-loxC5-SV40. Statistical t tests show ** = $p < 0.01$, * = $p < 0.05$., and n.s.

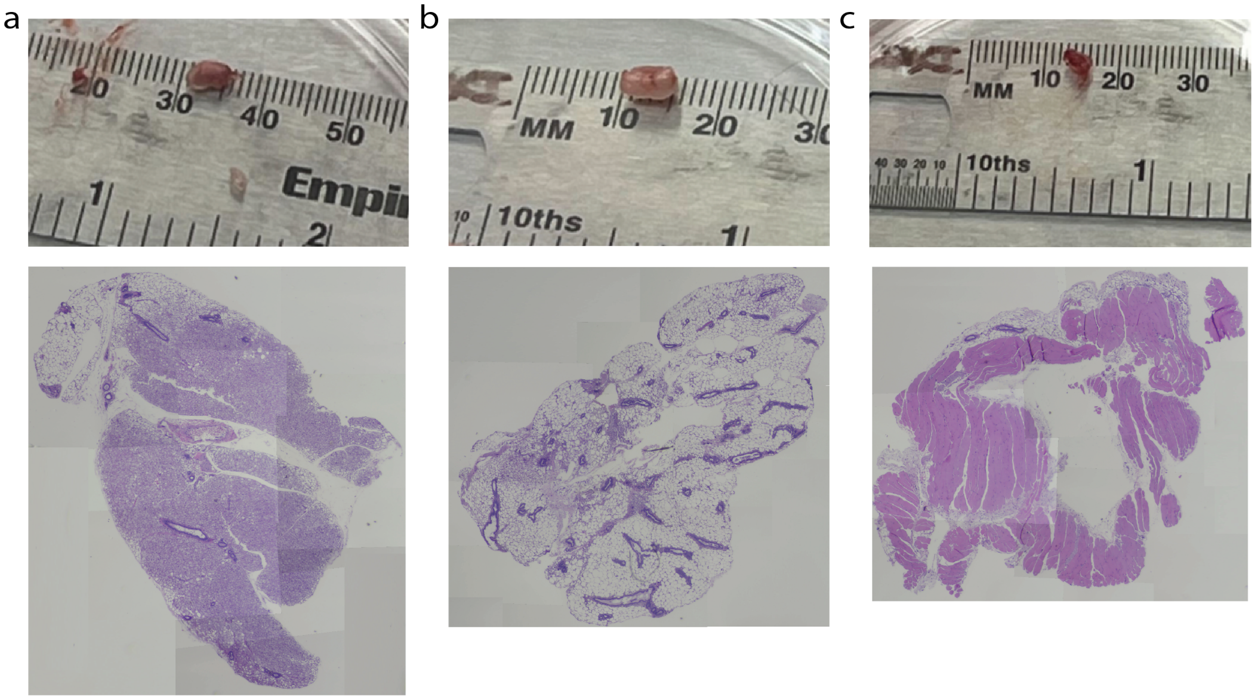


**Supplementary Figure 16: Teratomas.** Tumors extracted from hind legs of immune-compromised mice post-emiPSC injection (TOP) and H&E cross-sections (BOTTOM). All tumors surgically removed after 5.5 weeks and measured as pictured. **a.** Tumor from C1-loxC4-TP53shRNA2 **b.** Tumor from C1-loxC5-SV40 **c.** Tumor from C1-loxC6-SV40.

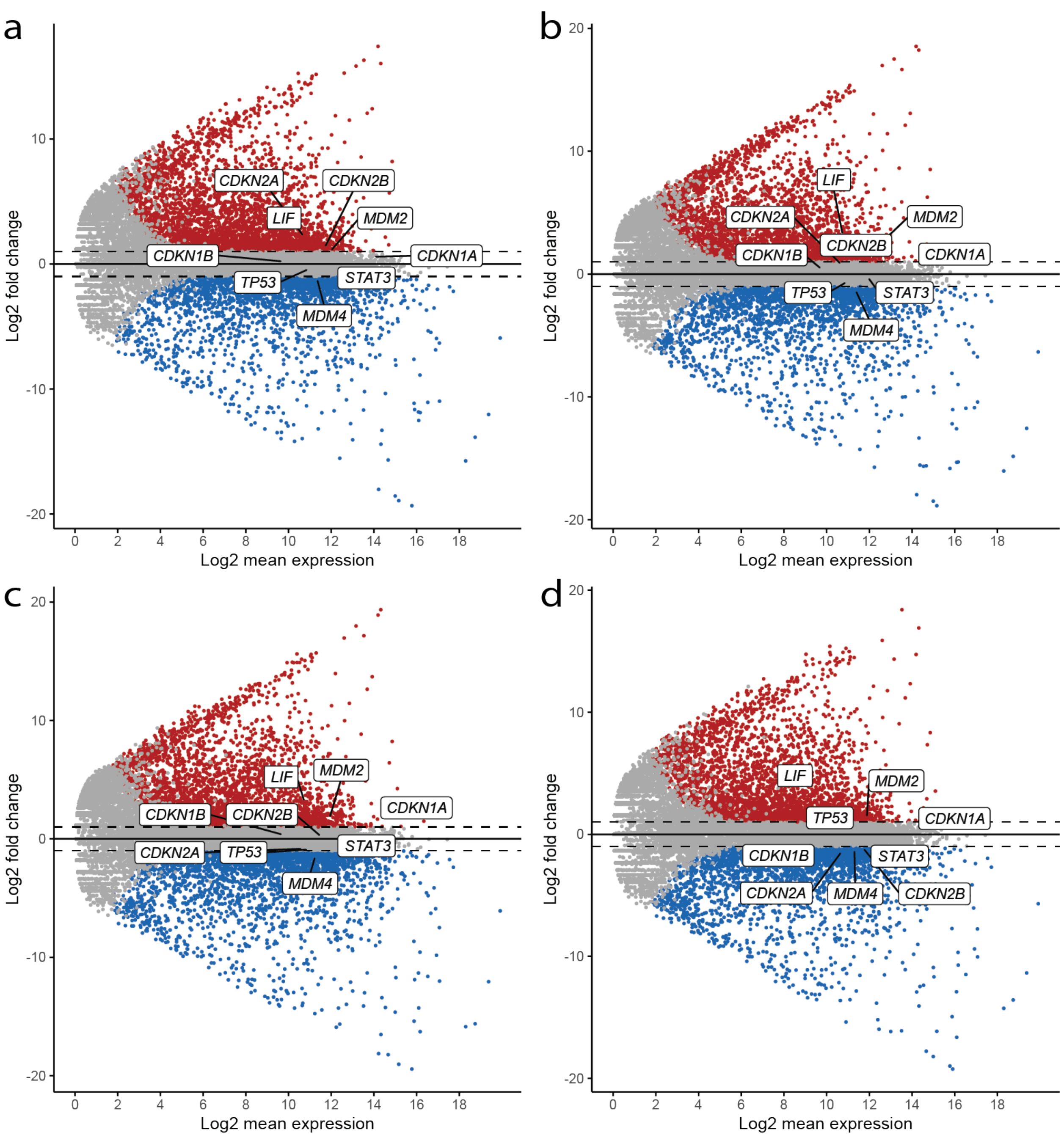


**Supplementary Figure 17: MA plots for emiPSCs.** MA plots highlighting core regulatory genes that control cell growth rate that are often strongly down-regulated in iPSCs from other species. **a.** C1-loxC4-SV40. **b.** C1-loxC5-SV40. **c.** C1-loxC6-SV40. **d.** C1-loxC4-TP53shRNA2.

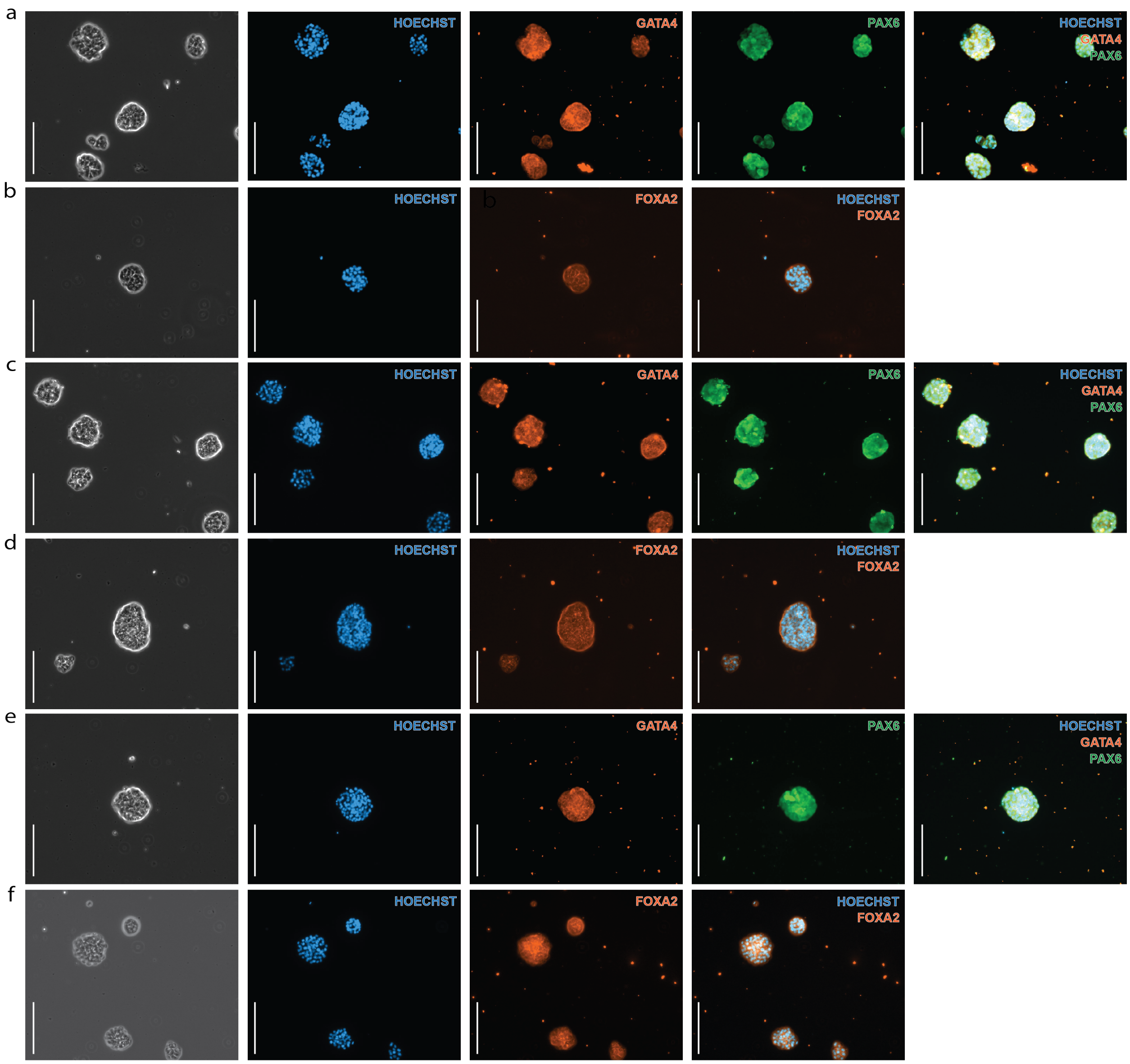


**Supplementary Figure 18: Embryoid bodies for cell additional cell lines stained for early differentiation markers.** Immunofluorescence (IF) microscopy images of embryoid bodies (EBs) formed by emiPSC lines. All cells shown at 10X magnification, scale bar = 200 $\mu$M **a.** C1-loxC4-SV40 stained for *PAX6* (ectoderm) and *GATA4* (mesoderm). **b.** C1-loxC4-SV40 stained for *FOXA2* (endoderm). **c.** C1-loxC6-SV40 stained for *PAX6* (ectoderm) and *GATA4* (mesoderm). **d.** C1-loxC6-SV40 stained for *FOXA2* (endoderm). **e.** C1-loxC4-TP53shRNA2 stained for *PAX6* (ectoderm) and *GATA4* (mesoderm). **f.** C1-loxC4-TP53shRNA2 stained for *FOXA2* (endoderm).
